# The spatiotemporal distribution of LIN-5/NuMA regulates spindle orientation and tissue organization in the *C. elegans* germ line

**DOI:** 10.1101/2024.08.31.610619

**Authors:** Réda M. Zellag, Vincent Poupart, Takefumi Negishi, Jean-Claude Labbé, Abigail R. Gerhold

## Abstract

Mitotic spindle orientation sets the cell division plane and is thus critical for maintaining tissue organization. The *C. elegans* gonad is tube-shaped, with germ cells forming a circumferential monolayer around a shared inner core of cytoplasm called the rachis. Each germ cell is connected to the rachis via a stable cytoplasmic bridge, polarizing germ cells along their rachis-basal axis. How this tissue organization is maintained during development is unclear, as germ cells lack the canonical cell-cell junctions that, in other tissue types, ensure proper spindle orientation. Here we use live-cell imaging of *C. elegans* germ cells, both *in situ* and in gonad explants, to show that the microtubule force generator dynein and its conserved regulator LIN-5/NuMA regulate spindle orientation in *C. elegans* germ cells and are required for germline tissue organization. We uncover a cyclic, polarized pattern of LIN-5/NuMA cortical localization that predicts centriole/centrosome positioning throughout the cell cycle, providing a means to align spindle orientation with the tissue plane. This work reveals a new mechanism by which oriented cell division can be achieved to maintain tissue organization during animal development.

## Introduction

The ability of organs to perform specialized functions such as nutrient absorption or reproduction depends upon the acquisition of proper tissue architecture during development. This requires the coordination of individual cell behaviours, including the orientation of cell division, which, by determining the position of daughter cells within the developing tissue, can impact both tissue shape and organization [1, 2]. Oriented cell division relies on cortical force generators, typically the molecular motor protein dynein and its activator <u>nu</u>clear <u>m</u>itotic <u>a</u>pparatus (NuMA), that act on astral microtubules to position the mitotic spindle and thereby dictate the axis of cell division [3].

Several mechanisms have been described by which the activity and/or localization of cortical force generators is regulated to generate oriented cell divisions. For example, spindles in epithelial cells are typically oriented perpendicular to the apical-basal cell axis, and thus parallel to plane of the tissue, and can be influenced by tissue-scale tension or patterning to provide an additional orientation bias within the plane of the tissue (so-called planar orientation). Apical-basal orientation involves concentrating the force generating machinery along lateral cell cortices [4, 5], by coupling its localization to apical-basal polarity and/or cell-cell junctional cues [6–12]. Planar orientation has been linked to the often interdependent phenomena of tissue tension [13–18], cell shape [16–22] and planar cell polarity [1, 23–25]. In most cases, planar orientation is also dictated by the association of cortical force generators with cell junctions (e.g. tri-cellular junctions [20, 21] and/or cell adhesion proteins (e.g. Cadherins [15]). However, several exceptions have been noted [13, 15], and examples of novel mechanisms are still emerging (e.g. [26]). Thus, a full picture of the diversity of mechanisms regulating spindle orientation during development is lacking.

The *C. elegans* gonad is a well-established model for the study of germline tissue development, yet relatively little is known about how germ cell spindle orientation is regulated. In adult hermaphrodites, the gonad is arranged into two symmetric U-shaped arms, each capped at its distal end by a Distal Tip Cell (DTC) that serves as a niche for the underlying mitotic germ cells [27, 28]. Within each gonad arm, germ cells are arranged in a rough circumferential monolayer around a common, central core of cytoplasm called the rachis (**Figure 1A**, [29, 30]). Each germ cell is connected to the rachis via a single cytoplasmic bridge that is maintained by a stable actomyosin ring, polarizing germ cells along their rachis-basal axis and forming a tissue-scale, tensile actomyosin network at the rachis surface [30–33]. Germ cell cytoplasmic bridges constrict in mitosis, but rachis markers remain enriched on their rachis face (**Figure 1B**, [34]). This architecture is present throughout development [31, 35], as the number of germ cells increases from two primordial germ cells in newly-hatched L1 larvae to ∼2000 germ cells in adults [27, 28], and as the gonad elongates along its distal/proximal (D/P) axis, undergoing a 180° turn to attain its final U-like shape [27, 36].

**Figure 1.**
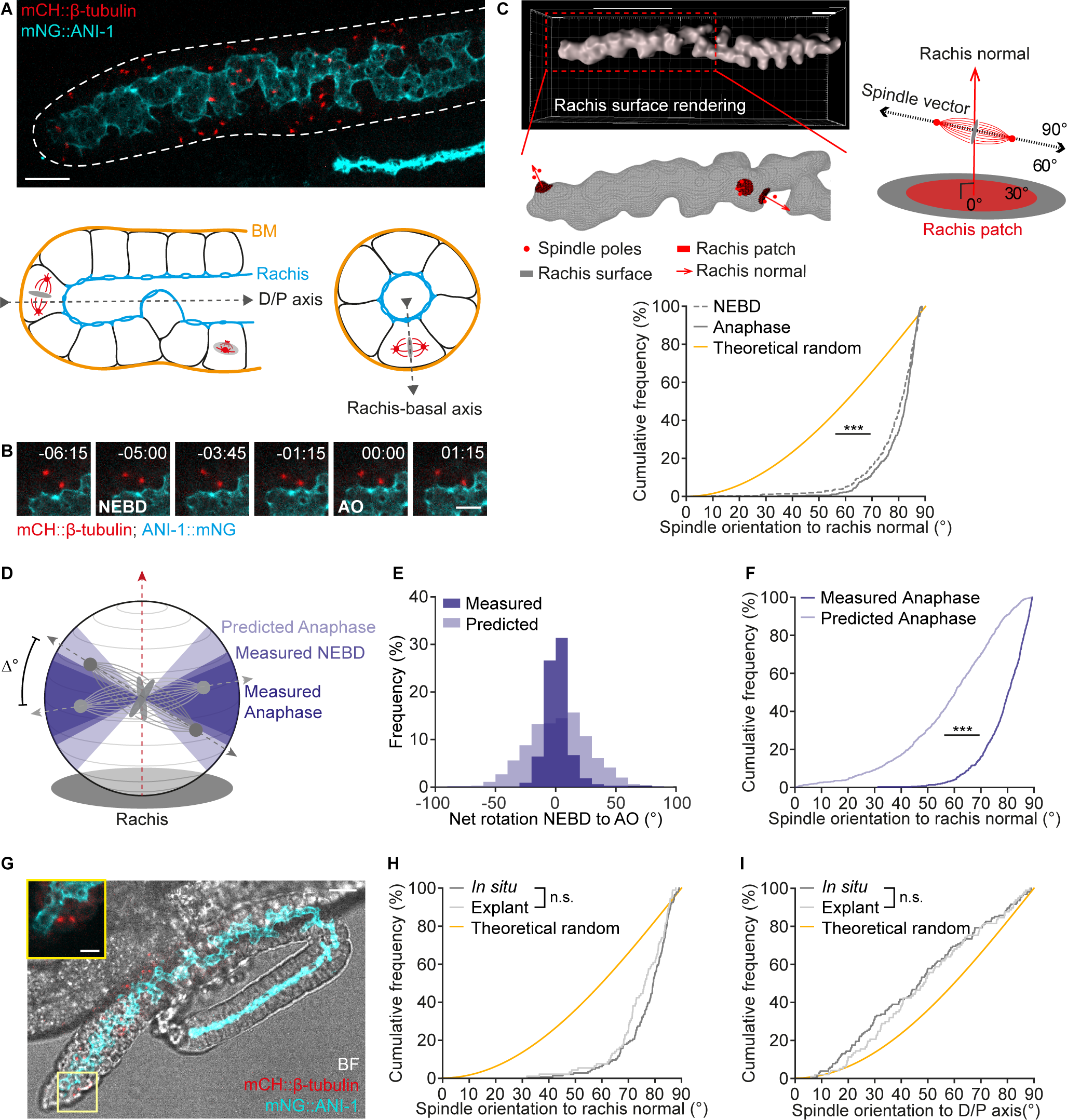
Germ cell spindles orient parallel to the rachis surface *in situ* and in gonad explants. (A) Top: Maximum intensity projection of a gonad arm from an L4 larva expressing mCH::β-tubulin (red) in the germ line and endogenously tagged mNG::ANI-1 (cyan) to mark spindles and the rachis, respectively. Scale bar = 10 µm. Bottom: Schematic representations of a distal gonad arm viewed length-wise (left) and *en face* (right). BM = basement membrane, D/P = distal/proximal. (B) Maximum intensity projections of a germ cell, expressing mCH::β-tubulin (red) and mNG::ANI-1 (cyan), undergoing mitosis. Numbers indicate time in minutes relative to anaphase onset. NEBD = nuclear envelope breakdown, AO = anaphase onset. Scale bar = 5 µm. (C) Representative rachis surface rendering from the mNG::ANI-1 signal (top) with a reconstruction of the rachis surface, showing a subset of mitotic spindles in relation to their respective rachis surface patches and rachis normal vectors (bottom). Scale bar = 10 μm. A schematic depicting spindle orientation to the rachis surface as the angle formed between the spindle vector and the rachis normal vector is shown on the right. Bottom: Cumulative distribution of spindle angles to the rachis normal for germ cells in L4 larvae at NEBD and anaphase as compared to the theoretical random distribution (yellow). (D) Schematic representation of the range of measured spindle angles to the rachis normal at NEBD and anaphase, as compared to the predicted range at anaphase (see Methods). Δ° represents the net rotation of a spindle, NEBD to AO. (E) Histogram comparing the measured versus predicted net spindle rotation between NEBD and AO. The observed range of rotation is narrower than what is predicted by measured spindle dynamics. (F) Cumulative distribution of the measured versus predicted spindle angles to the rachis normal at AO, showing a significant bias in measured angles. Measured data are reproduced from (C). (G) Bright field image of a gonad explant from an L4 larva, overlaid with a maximum intensity projection of mNG::ANI-1 (cyan) and mCH::β-tubulin (red). Scale bar = 10 μm. Inset shows mNG::ANI-1 (cyan) and mCH::β-tubulin (red) in a mitotic germ cell. Scale bar = 5 μm. (H-I) Cumulative distribution of spindle angles to the rachis normal (H) and the D/P axis (I) for measurements made at anaphase *in situ* and in explants, with the theoretical random distribution shown for comparison. *In situ* data were drawn at random from the dataset used in Figure S1B. For all panels, n.s. = *p* > 0.05, *** = *p* < 0.001. Summary statistics and statistical tests used are given in Table S4. See also Figures S1 and S2 and Video S1.

*C. elegans* germ cells, like epithelial cells, are thought to divide within the plane of the germline tissue (**Figure 1A-B**, [34]). They also exhibit a mild orientation bias relative to the gonadal D/P axis [37]. The former could serve to maintain the germline monolayer and ensure that each daughter cell remains connected to the rachis after cell division, while the latter could contribute to, or arise from, the elongated gonad tissue shape. However, the mechanisms aligning germ cell spindles to either axis are unknown and the functional consequences of disrupting spindle orientation have not been assessed. Moreover, electron microscopy analyses suggest that *C. elegans* germ cells lack cell-cell junctions [29, 30] and are therefore without a conventional means for orienting cell division.

Here we use live-cell imaging of *C. elegans* germ cells *in situ* and in gonad explants to investigate the regulation of spindle orientation during gonad development. We find that germ cell spindles are oriented parallel to the rachis surface and that spindle orientation to both the rachis surface and gonadal D/P axis occurs independently of interphase cell shape and forces external to the gonad. Instead, both spindle orientation biases require the force generators dynein and LIN-5/NuMA and depletion of either regulator leads to disorganization of the germline tissue. By characterizing the cortical localization of LIN-5/NuMA and centriole/centrosome positioning in interphase and mitotic germ cells, we propose a model wherein the exclusion of LIN-5/NuMA from the germ cell rachis surface throughout the cell cycle, coupled with the basal localization of centrioles during interphase, establishes spindle orientation parallel to the rachis surface during prophase. The dynamic association of LIN-5/NuMA with lateral cortices adjacent to spindle poles during mitosis then maintains this orientation into anaphase, thus positioning both daughter cells in the plane of the tissue and maintaining germline tissue organization.

## Results

### Mitotic spindles are oriented parallel to the rachis surface in *C. elegans* germ cells

To accurately assess spindle orientation in dividing *C. elegans* germ cells, we first developed an approach to identify the rachis surface for each cell and to measure spindle orientation relative to this surface through time. This task was complicated by the fact that the rachis core is tortuous [34] and that germ cells can divide within the 3D tube-like structure of the gonad at any angle relative to the imaging plane (see **Figure 1A-B and Video S1**). We used animals that express fluorescent protein (FP)-tagged versions of β-tubulin to track and pair centrosomes [37, 38] and the actomyosin scaffold protein anillin (ANI-1 [31, 39]) to generate a 3D rendering of the rachis surface (**Figure 1C**). We then identified the rachis patch nearest to each mitotic spindle and represented its orientation as the normal vector to the rendered surface (hereafter the rachis normal). Spindle orientation in relation to the rachis surface was then defined as the angle formed between the spindle vector and the rachis normal vector in 3D (**Figure 1C**; see Methods). Accordingly, spindles perfectly parallel to their relevant rachis surface will be orthogonal to the rachis normal, with a spindle angle of 90°. Measurements were done on L4 larvae, a developmental stage characterized by robust germ cell proliferation and sustained gonad growth [27, 28].

We found that the majority of germ cell spindles are roughly orthogonal to the rachis normal in anaphase (and thus parallel to the rachis surface), with 98% of spindles having an angle of 60° or greater (**Figure 1C**). We calculated the theoretical range of angles that spindles could assume during anaphase elongation without forcing spindle poles into contact with the rachis surface plane (see Methods) and found a good concordance with our measured values – in theory, an angle of 53° or greater, relative to the rachis normal, should prevent spindle poles from contacting the rachis surface and, across all cells, only 2 of 503 spindles had a measured angle less than this value (**Figure S1A-B**). Thus, the range of spindle angles that we observe is consistent with a relative lack of spindle pole engagement with the rachis surface.

The spindle orientation bias relative to the rachis surface is stronger than the bias that we previously reported relative to the gonadal D/P axis [37]. However, like the bias to the D/P axis, spindle orientation parallel to the rachis surface was set up during prophase, with a similar distribution of spindle angles at nuclear envelop breakdown (NEBD) as in anaphase (**Figure 1C)**. We considered the possibility that a bias in spindle orientation at NEBD would be sufficient to ensure that spindles maintain this orientation into anaphase. We measured the magnitude of spindle rotations relative to the rachis normal during prometa/metaphase and used this to predict the range of angles that would be observed at anaphase onset if spindles were not constrained (**Figure 1D**). We found that spindles explore a comparatively narrow range of orientations during prometa/metaphase and that without an active mechanism to maintain the orientation bias to the rachis surface, spindle orientation in anaphase would be considerably more variable (**Figure 1E-F**). Taken together, we conclude that germ cells divide parallel to their rachis surface, and thus within the plane of the gonad tissue. Spindle orientation relative to the rachis surface is established during prophase and spindle rotations during prometa/metaphase tend to be oscillatory such this orientation bias is maintained through anaphase.

### Germ cell spindles orient parallel to the rachis surface irrespective of developmental stage and distance to the niche

We next assessed whether spindle orientation was influenced by developmental changes in tissue organization. Previous work has suggested that the rachis becomes increasingly tortuous, with the germ line adopting a folded, epithelia-like organization as animals progress from the L4 larval stage into adulthood (**Figure S1C**, [34]). We found that germ cell spindles were oriented parallel to the rachis surface both in L3 larvae, when the germ line lacks folds, and in 1-day old adults, in which germ line folds are more pronounced (**Figure S1C-D**).

We also considered whether germ cell position along the gonadal D/P axis, and thus distance from the niche, affected spindle orientation (**Figure S1E**). However, the same spindle orientation bias was present in all regions of the distal gonad, from the distal-most region to the mitotic-to-meiotic transition zone (**Figure S1F**). Together, these results indicate that spindle orientation relative to the rachis surface is robust to changes in germline organization during development and germ cell differentiation as cells exit the niche.

### Spindle orientation in germ cells is gonad-autonomous

The gonad in L4 larvae consists of the germ line and somatic gonadal cells, including the DTC niche and the gonadal sheath cells which enwrap each gonad arm, and the entire organ is radially constrained by the shape of the animal and adjacent tissues [27]. To determine whether spindle orientation in germ cells is influenced by mechanical constraints external to the gonad, we tracked germ cell divisions in gonad explants. We extruded gonads into a medium previously shown to be permissible for *C. elegans* embryonic blastomere culture [40] (**Figure 1G**; see Methods). Following extrusion, explants retained somatic gonadal cells and the gonadal basement membrane (**Figure S2A-B**) and maintained proper germline architecture (**Figure 1G**), suggesting that they are largely intact. We found that mitotic duration and the rate of spindle elongation in germ cells dividing within these explants were similar to what we observed *in situ* (**Figure S2C-D**), indicating that this explant culture approach is physiologically relevant. Interestingly, germ cells in explants showed similar spindle orientation biases to cells *in situ*: spindle orientation in anaphase was strongly biased parallel to the surface of the rachis and weakly biased relative to the D/P axis of the tissue (**Figure 1H-I**). These results indicate that spindle orientation in germ cells is independent of constraints imposed by other anatomical features of the animal and that the mechanisms regulating spindle orientation are autonomous to the gonad tissue.

### The cell long axis does not predict spindle orientation in germ cells

In many cell types, the cell’s long axis predicts the orientation of division (the so-called ‘long axis’ rule, as proposed by Hertwig [41] and observed in several instances, e.g. [17, 19, 20, 22, 42, 43]). Thus, spindle orientation in germ cells could be a consequence of germ cell shape. To address this possibility, we assessed germ cell shape in mitotic and interphase cells using animals in which centrosomes (β-tubulin), the rachis surface (either ANI-1 or the septin UNC-59 [37, 44]) and the plasma membrane (the PH domain of rat PLC1∂ [45, 46]) were marked (**Figure 2A**). Rendering of the membrane marker in 3D allowed us to fit an ellipsoid to each germ cell and determine the length and orientation of the cell’s major (i.e. long) and minor axes. We found that germ cells were mildly anisotropic, with mitotic cells being on average larger and more spherical than the largest interphase cells (**Figure 2B-C**), consistent with cell growth occurring prior to mitotic entry and mitotic cell rounding.

**Figure 2.**
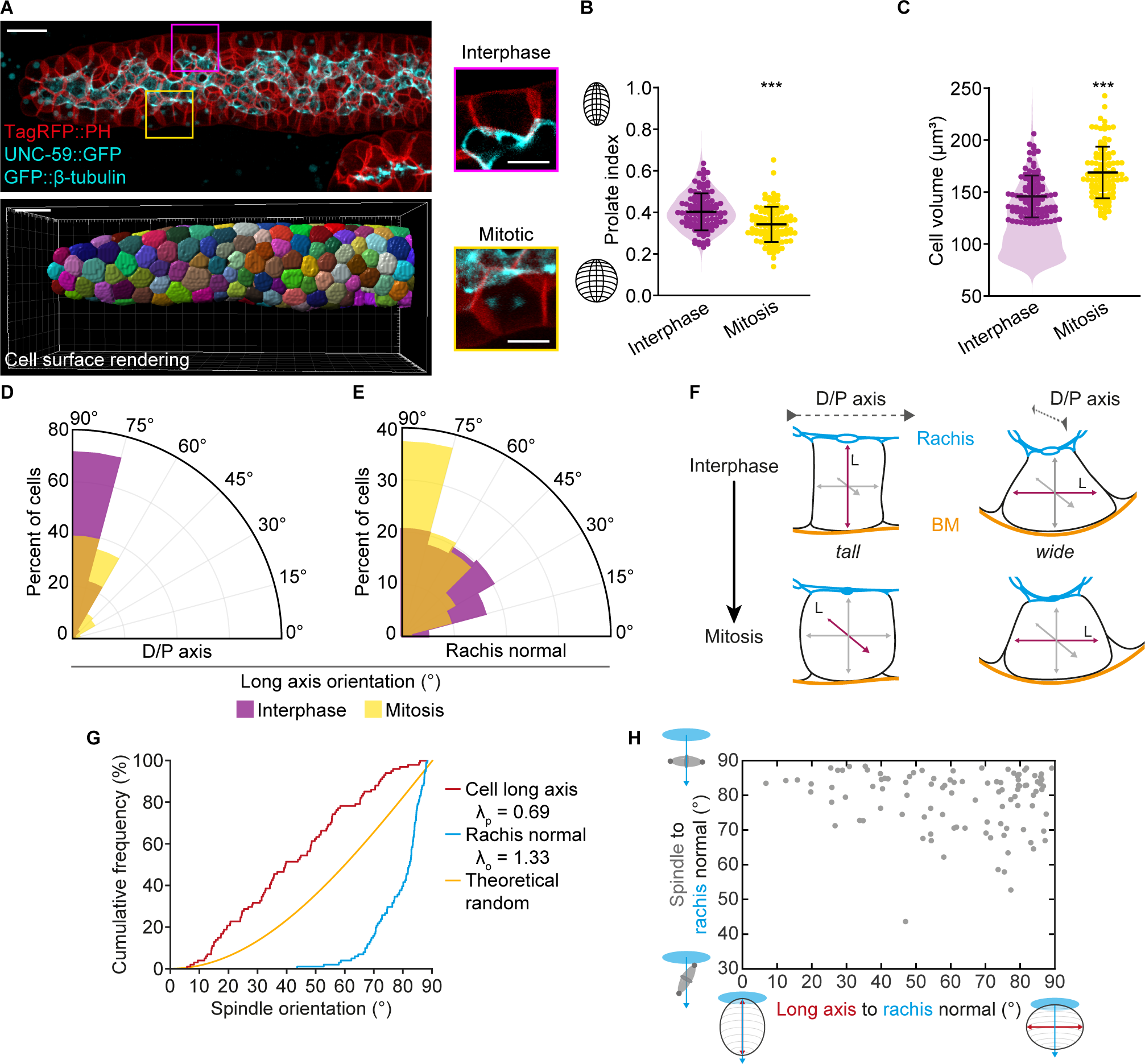
The cell long axis does not predict spindle orientation in germ cells. (A) Maximum intensity projection (top) and 3D cell surface rendering (bottom) of a gonad arm from an L4 larva expressing endogenously tagged UNC-59::GFP to mark the rachis (cyan), and GFP:: β-tubulin and TagRFP::PH in the germ line, to mark spindles (cyan) and cell membranes (red), respectively. Scale bar = 10 μm. An interphase (magenta boxed) and mitotic (yellow boxed) germ cell are shown to the right. Scale bar = 5 μm. (B-C) The prolate index (B) and cell volume (C) of germ cells in interphase and mitosis. For interphase cells, violin plots show the data for all cells and dots show the data used for comparison with mitotic cells. Interphase cells were selected based on cell volume by drawing a random set of cells from all interphase cells with a cell volume within ± 2 standard deviations of the mean volume for mitotic cells. Each dot represents one cell. Bars represent the mean ± standard deviation. *** = *p* < 0.001 (D-E) Rosette plots showing the orientation of the cell long axis in interphase (magenta) and mitosis (yellow) to the D/P axis (D) and rachis normal (E). (F) Schematic representation of interphase and mitotic germ cell shape, based on the measurements in (D) and (E), depicting configurations of the cell long axis orientation to the D/P axis and rachis surface. (G) Cumulative distribution of spindle angles to the cell long axis in mitosis as compared to the rachis normal for the same set of cells. Spindles show a stronger bias (λ_o_ = 1.33) orthogonal to the rachis normal than parallel (λ_p_ = 0.69) to the cell long axis. (H) Scatter plot showing the relationship between spindle orientation to the rachis normal relative to the angle between each cell’s long axis and the rachis normal, where an angle of 0° represents perfect parallel alignment. Summary statistics and statistical tests used are given in Table S4.

To determine whether the interphase cell long axis predicted spindle orientation in mitosis, we assessed its orientation to both the rachis normal and the D/P axis. We found that the interphase cell long axis could adopt roughly any orientation with respect to the rachis normal but was skewed orthogonal to the D/P axis (**Figure 2D-E**). This suggests that germ cell shape is slightly compressed relative to the gonadal D/P axis, with cell volume redistributing either as increased cell “height” along the rachis-basal axis or cell “width” orthogonal to D/P axis (**Figure 2F**). Importantly, these results show that the cell long axis in interphase does not align well with the orientation of germ cell division, particularly with respect to the rachis surface.

We next looked at the relationship between spindle orientation and the cell long axis during mitosis, as in several other models spindles can also respond to mitotic cell shape [16, 42, 43, 47, 48]. In mitotic cells, the orientation of the cell long axis was more likely to be parallel to the rachis surface and less likely to be perpendicular to the D/P axis than in interphase cells (**Figure 2D-E)**, suggesting that cell rounding occurs primarily orthogonal to the rachis normal and in the plane of the tissue (**Figure 2F**). We found that spindles tended to align with the cell long axis at anaphase, but that this bias was relatively weak compared to the bias in spindle orientation to the rachis normal for the same cells (**Figure 2G**). We then asked whether the orientation of the cell long axis in mitosis influenced spindle orientation relative to the rachis normal. However, even in cells where the long axis and rachis normal were relatively well aligned, spindles were still oriented orthogonal to the rachis (**Figure 2H**). Together, these results show that interphase cell shape does not predict spindle orientation in *C. elegans* germ cells, and that in mitotic cells, the rachis surface is the primary determinant of germ cell spindle orientation.

### Dynein and LIN-5/NuMA are required for germ cell spindle orientation and germline tissue organization

Dynein and LIN-5/NuMA regulate spindle orientation in the *C. elegans* one-cell zygote [49], but a role for these proteins in germ cells has not been assessed. As both regulators are required for numerous cellular processes throughout development, we used the <u>a</u>uxin-inducible <u>d</u>egron (AID) system[50, 51] to acutely deplete dynein heavy chain (DHC-1) and LIN-5/NuMA specifically in the germ line. For technical reasons, we used the gonad surface as a proxy for the rachis surface. Treating L4 larvae with auxin for 40 minutes reduced germline DHC-1/dynein and LIN- 5/NuMA to roughly 40% and 20% of their control levels, respectively (**Figures 3A-B**) and significantly decreased spindle movement during prometa/metaphase and spindle elongation in anaphase, consistent with a substantial loss of cortical pulling forces (**Figures S3A-C**). Under both conditions, germ cell spindle orientation in anaphase was strongly perturbed: the spindle orientation bias relative to the gonad surface was significantly reduced and spindle orientation relative to the D/P axis was effectively randomized (**Figure 3C-D**).

**Figure 3.**
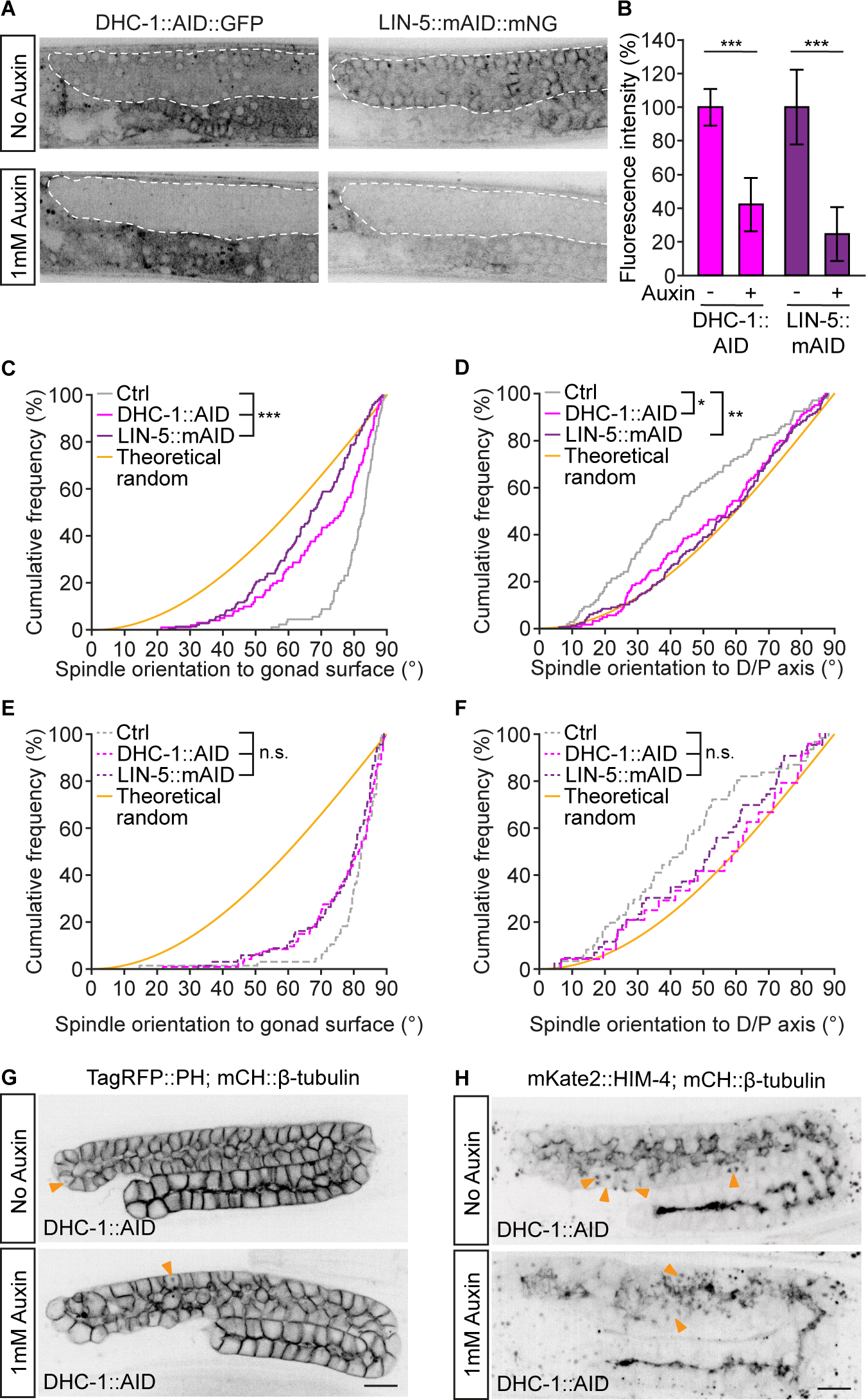
Dynein and LIN-5/NuMA are required for germ cell spindle orientation and germline tissue organization. (A) Confocal sections through the mid-plane of gonad arms from L4 larvae expressing DHC-1::AID::GFP (dynein, left) or LIN-5::mAID::mNG (NuMA, right), with a germline-specific TIR1 (*sun-1p*::*TIR1*), after a 40-minute auxin treatment, as compared to untreated animals. The germ line is outlined in white. Scale bar = 10 μm. (B) Quantification of depletion by fluorescence intensity measurements of each GFP/mNG-tagged protein in the germ line. Measurements were normalized to the mean value for non-depleted germ lines. *** = *p* < 0.001 (C-F) Cumulative distributions of spindle angles to the gonad surface normal vector (C, E) and the D/P axis (D, F) for measurements made at anaphase (C-D) and NEBD (E-F) comparing control animals to those depleted of DHC-1/dynein or LIN-5/NuMA as in (A-B). Depletion of either DHC-1/dynein or LIN-5/NuMA reduces the spindle orientation bias to the gonad surface and D/P axis. The theoretical random distribution (yellow) is shown for reference. n.s. = *p* > 0.05, * = *p* < 0.05, ** = *p* < 0.01, *** = *p* < 0.001 (E) Single confocal sections of gonad arms from L4 larvae expressing mCH::β-tubulin and TagRFP::PH in the germ line, depleted (bottom) or not (top) of DHC-1/dynein in the germ line by a 6-hour auxin treatment. (F) Maximum intensity projections of gonad arms from L4 larvae expressing mCH::β-tubulin in the germ line and endogenously tagged mKate2::HIM-4 to mark the rachis, depleted (bottom) or not (top) of DHC-1/dynein in the germ line by a 6-hour auxin treatment. Yellow arrows indicate spindles. Scale bars = 10 μm. Summary statistics and statistical tests used are given in Table S4. See also Figure S3.

Given the average duration of a germ cell mitosis (30 minutes, prophase through anaphase [52]), most cells entering anaphase after 40 minutes of auxin treatment likely experienced DHC- 1/dynein or LIN-5/NuMA depletion starting in prophase. We therefore asked whether spindle orientation at NEBD was also affected and found that indeed, in the subset of cells where both NEBD and anaphase occurred during our imaging window, both spindle orientation biases were reduced at NEBD, albeit to a lesser extent compared to anaphase (**Figure 3E-F**). These results indicate that DHC-1/dynein and LIN-5/NuMA are required to both set up proper spindle orientation at NEBD and to maintain it during mitosis.

We next asked whether depleting DHC-1/dynein or LIN-5/NuMA affected germline tissue organization by subjecting animals to auxin treatment for 6 hours, during which most mitotically competent cells will have undergone a single round of division [27, 53]. We found that a 6-hour depletion of either protein resulted in germline disorganization, with a notable alteration of rachis shape and increased variation in germ cell size (**Figures 3G-H and S3D**). Together, these results indicate that DHC-1/dynein and LIN-5/NuMA are important for both proper germline tissue organization and germ cell spindle orientation, consistent with a causal link between the orientation of germ cell divisions and the maintenance of germline architecture during development.

### The cortical distribution of LIN-5/NuMA predicts germ cell spindle orientation

Since LIN-5/NuMA is required for proper spindle orientation in germ cells, and its cortical localization correlates with sites of force generation in other cell types [3, 54], we examined how its cortical localization in germ cells related to germ cell spindle orientation biases. We measured the fluorescence intensity of FP-tagged LIN-5/NuMA at the rachis, basal and lateral cell cortices, in both interphase and mitotic germ cells, using a custom 3D line-scan method and normalizing the intensity of FP-tagged LIN-5/NuMA to the membrane signal at each cell cortex (**Figure 4A-C and S4A-C**).

**Figure 4.**
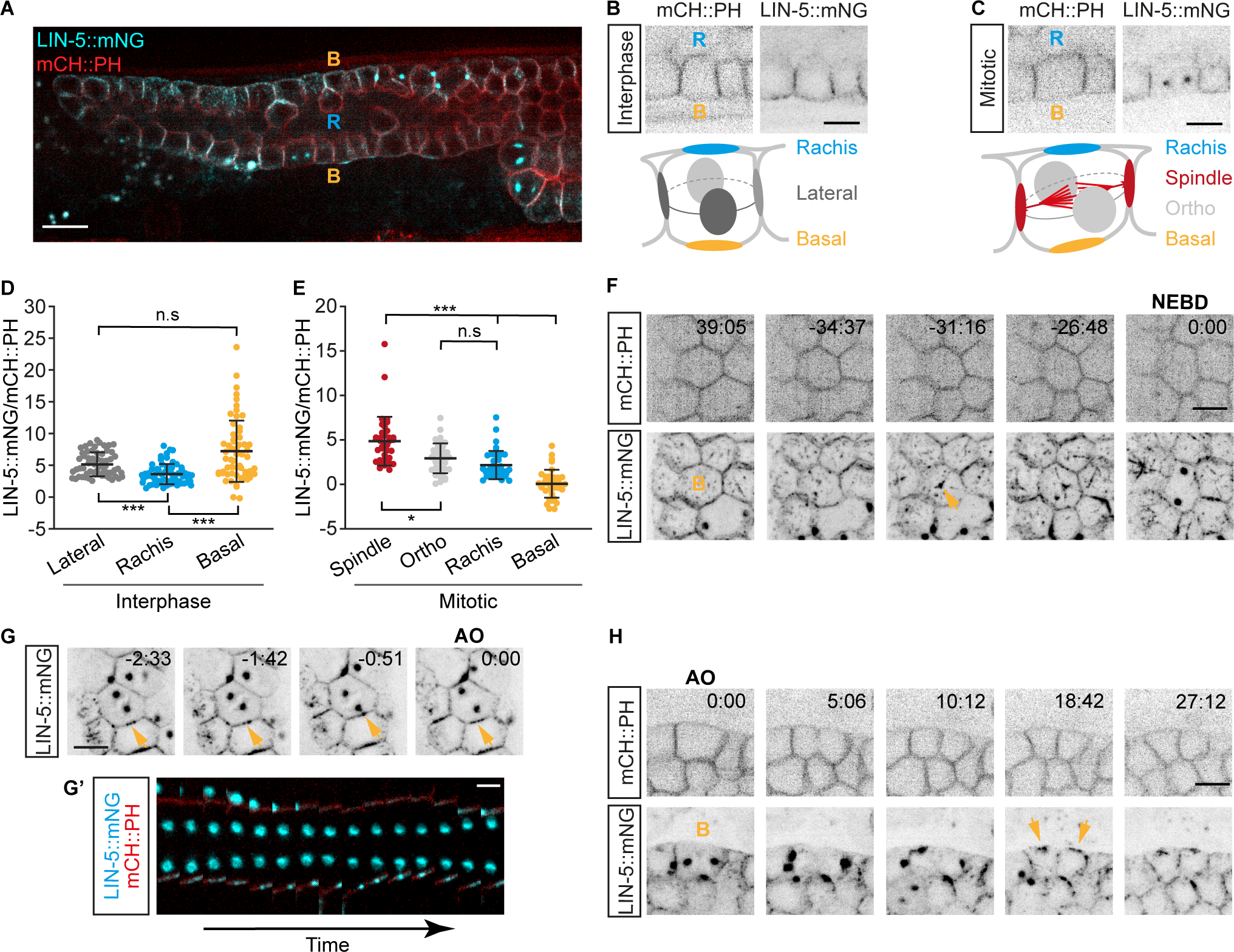
The cortical distribution of LIN-5/NuMA predicts germ cell spindle orientation. (A) Maximum intensity projection of confocal sections through the middle of a gonad arm from an L4 larva expressing endogenously tagged LIN-5::mNG (NuMA, cyan) and germline-expressed mCH::PH to mark cell membranes (red). B = basal, R = rachis. Scale bar =10 μm. (B-C) Maximum intensity projections (top) of germ cells during interphase (B) and mitosis (C) showing cell membranes (mCH::PH; left) and LIN-5::mNG (right) in inverted greyscale. Scale bars = 5 μm. Schematic representations (bottom) depict the line scan method used for fluorescence intensity measurements at cell cortices along the rachis-basal and spindle/orthogonal or lateral axes. (D-E) LIN-5::mNG fluorescence intensity, normalized to the membrane mCH::PH signal, at the indicated interphase (D) and mitotic (E) germ cell cortices, as depicted in (B) and (C). Dots represent per cell measurements. Bars represent the mean ± standard deviation. n.s. = *p* > 0.05, * = *p* < 0.05, *** = *p* < 0.001 (F) Maximum intensity projections of the mCH::PH (top) and LIN-5::mNG (bottom) signal at the basal surface (top-down view) of a germ cell progressing through prophase. Time is shown in minutes relative to nuclear envelop breakdown (NEBD). B = basal and marks the cell of interest. Arrow head indicates the appearance of a LIN-5::mNG puncta at the basal surface. Scale bar = 5 µm. (G) Maximum intensity projections of the LIN-5::mNG signal at the mid-plane of a germ cell in mitosis. Time is shown in minutes relative to anaphase onset (AO). Arrow heads indicate LIN-5::mNG at the lateral cell surface adjacent to the spindle pole. Scale bar = 5 µm. (G’) Kymograph along the spindle vector for the cell shown in (G) with mCH::PH (red) and LIN-5::mNG (cyan) showing the formation and dissolution of a LIN-5::mNG focus (lower membrane; bright spot adjacent to upper membrane is the spindle pole from a neighboring cell). Time scale bar = 17 seconds. (H) Maximum intensity projections of the mCH::PH (top) and LIN-5::mNG (bottom) signal at the mid-plane of a germ cell in anaphase showing a lateral view of the basal surface (B) and the basal localization of LIN-5::mNG upon mitotic exit (arrow heads). Time is shown in minutes relative to anaphase onset (AO). Scale bar = 5 µm. Summary statistics and statistical tests used are given in Table S4. See also Figure S4 and Videos S2-S4.

In interphase cells, LIN-5/NuMA was enriched on lateral and basal cortices, while its levels were low at the rachis surface (**Figure 4D**). In mitotic cells, LIN-5/NuMA underwent a pronounced change in its cortical association: while levels at the rachis surface remained low, the amount of LIN-5/NuMA on other cell cortices, particularly the basal cortex, was noticeably reduced (**Figure 4E**). Timelapse images showed a loss of LIN-5/NuMA from the basal surface during prophase that was concomitant with the appearance of basally located LIN-5/NuMA puncta, likely at the nascent centrosomes (**Figure 4F and Video S2**). During prometa/metaphase, LIN-5/NuMA localization was highly dynamic, appearing as puncta on the lateral cortices that tracked with centrosome movements (**Figure 4G and Video S3**). Correspondingly, at anaphase onset, we found that LIN-5/NuMA was enriched on lateral cortices along the spindle vector, and thus adjacent to centrosomes, relative to lateral cortices along the orthogonal vector (**Figure 4E**). In late anaphase/telophase, LIN-5/NuMA reappeared at the basal surface, seemingly in conjunction with the basal movement of the disassembling centrosomes (**Figure 4H and Video S4**).

Thus, during mitosis, cortical LIN-5/NuMA is dynamic, and its enrichment on lateral cell cortices along the spindle axis is consistent with its role in controlling the final orientation of cell division. Importantly, LIN-5/NuMA levels are consistently low at the germ cell’s rachis surface, suggesting a relative lack of pulling forces on astral microtubules from this cortex, both as spindle orientation is established in prophase and maintained during mitosis.

### Interphase centriole positioning and centrosome movement during prophase establish germ cell spindle orientation

As germ cells exit mitosis, LIN-5/NuMA reappears on the basal cortex coincident with the basal localization of LIN-5/NuMA foci, which are likely formed by the disassembling centrosomes (**Figure 4H and Video S4**). This suggests that centrosomes may be positioned at or near the basal germ cell cortex during interphase. To assess this, we used animals bearing FP-tagged ƴ-tubulin and histone H2B to visualize centriole position relative to germ cell nuclei in interphase cells. We found that centrioles were located close to the nucleus, towards the exterior of the gonad and thus the basal cell cortex (**Figure 5A**). We measured centriole position on the nucleus relative to the gonadal radial axis at each cell (here defined in 2D as the line passing from the center of the gonad, through the center of each nucleus in a cross-sectional view of the gonad; **Figure 5A**) and found that centrioles were indeed largely confined to positions along the basal 70% of the nuclear surface (**Figure 5B**). Thus, during interphase, centrioles are positioned on the basal face of the nucleus, roughly opposite to the rachis surface. This is distinct from the lateral position centrosomes occupy during anaphase, suggesting that they are repositioned upon mitotic exit.

**Figure 5.**
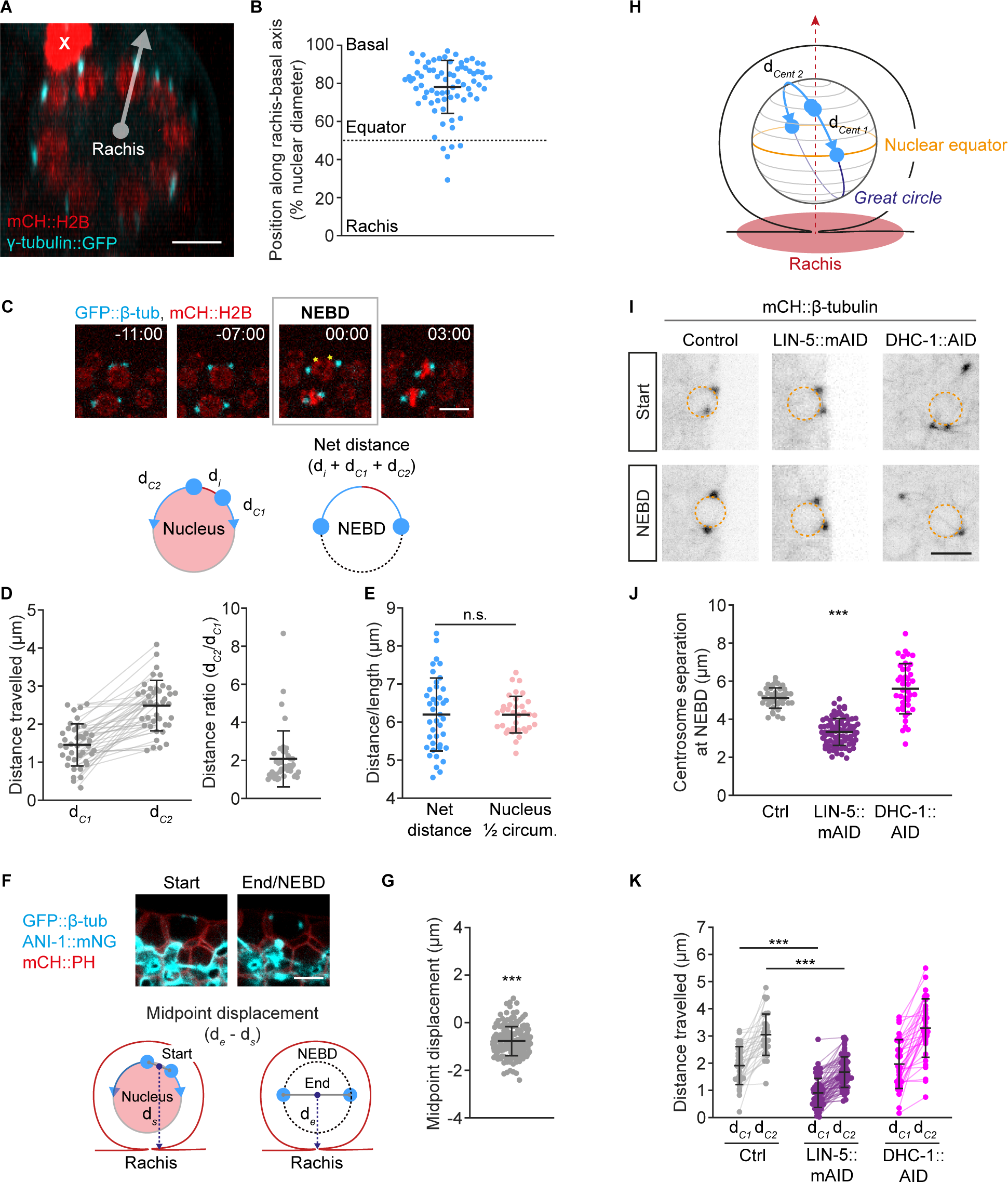
Interphase centriole positioning and centrosome movement during prophase establish germ cell spindle orientation. (A) Maximum intensity projection of a cross-sectional view of a distal gonad arm from an L4 larva expressing mCH::HIS-58(H2B) to mark nuclei (red) and TBG-1(ƴ-tubulin)::GFP to mark centrioles (cyan). Scale bar = 5 μm. The grey line depicts the rachis-basal axis for the nucleus at the top right. The bright red blob (X) is a somatic nucleus outside of the gonad. (B) The position of centrioles in interphase relative to the nuclear rachis- basal axis for each cell expressed as a percent of nuclear diameter with 100% and 0% indicating the basal and rachis poles, respectively. (C) Maximum intensity projections (top) of germ cells expressing mCH::HIS-11(H2B) (red) and GFP::β-tubulin (cyan). Time is shown in minutes relative to nuclear envelop breakdown (NEBD). Yellow asterisks represent centrosome position at the start of tracking. Scale bar = 5 μm. Schematics (below) depict centrosome migration (blue) around the nuclear surface (pink) during prophase, where the net distance travelled equals the sum of centrosome-to-centrosome distance at the start of tracking (d*_i_* ) plus the distance traveled by the two centrosomes prior to NEBD (d*_C1_* and d*_C2_*) centrosome. (D) The distanced travelled around the nucleus in prophase (left) with the distance ratio (d*_C2_*/d*_C1_*; right) for pairs of centrosomes showing that both centrosomes move, but one centrosome in each pair tends to move further. (E) Net distance traveled by centrosome pairs around the nucleus compared to the measured nucleus half circumference. (F) Maximum intensity projections (top) of a mitotic germ cell at the start of centrosome tracking in prophase and at NEBD with mNG::ANI-1 (cyan), GFP::β-tubulin (cyan) and TagRFP::PH (red) marking the rachis, spindle poles and cell membranes, respectively. Scale bar = 5 μm. Schematics (below) illustrate how centrosome-to-centrosome midpoint displacement relative to the rachis surface was measured. (G) Centrosome-to-centrosome midpoint displacement relative to the rachis surface from the start of centrosome tracking (d_s_) and end (d_e_) of prophase (NEBD). Displacement is significantly less than 0, indicating that, on average the centrosome-to-centrosome midpoint is closer to the rachis surface at NEBD. (H) Schematic representation of centrosome migration during prophase as inferred from the measurements in (A-G). Centrosomes travel along a nuclear “great circle” from a basal starting position to the nuclear equator parallel to the rachis surface. (I) Maximum intensity projections of mCH::β-tubulin in germ cells at the start of centrosome tracking (top) and at NEBD (bottom) in control animals (left) and animals depleted of LIN-5/NuMA (middle) or DHC-1/dynein (right) by a 40-minute auxin treatment. Scale bar = 5 μm. The nucleus is outlined in orange. (J-K) Centrosome-to-centrosome distance (separation) at NEBD (J) and the distanced travelled around the nucleus in prophase for pairs of centrosomes (K) in control animals and animals depleted of Lin-5/NuMA or DHC-1/dynein by a 40-min auxin treatment. Centrosome separation is reduced in LIN-5/NuMA depleted cells and more variable (*p* < 0.001, Bartlett test for equal variance) in DHC-1/dynein depleted cells. Centrosome migration in prophase is also reduced in LIN-5/NuMA depleted cells. In all panels, dots represent individual centrosomes (D and K) or per cell values (B, E, G and J), bars represent the mean ± standard deviation. n.s. = *p* > 0.05, *** = *p* < 0.001. Summary statistics and statistical tests used are given in Table S4. See also Figure S5.

The basal position of centrioles during interphase raised the question of how the spindle assembles parallel to the rachis surface in the following mitosis. We reasoned that if the two newly-duplicated centrosomes moved from the basal side of the nucleus towards the rachis surface during centrosome separation in prophase, this would bring them parallel to rachis surface by NEBD. To assess this, we tracked centrosomes throughout prophase and found that both centrosomes indeed migrated, with one centrosome typically travelling further than its partner (**Figure 5C-D**). After accounting for centrosome separation prior to the start of tracking (see **Figure 5C**), the combined distance travelled by pairs of centrosomes was roughly equivalent to half the nuclear circumference (**Figure 5E)**. To determine whether this movement was directed towards the rachis surface, we measured the displacement of the midpoint between centrosomes along the rachis-basal axis, and found that it moved, on average, closer to the cell’s rachis face (**Figure 5F-G)**. Together these results suggest that both centrosomes traverse the shortest possible route from a basal starting point, to reach opposite sides of the nucleus at the nuclear equator, effectively travelling along a nuclear “great circle”. This pattern of migration ensures that the spindle forms parallel to the rachis surface at NEBD (**Figure 5H**).

The directed movement of centrosomes requires a directional bias in the forces driving their migration [55]. Given the polarized cortical distribution of LIN-5/NuMA (**Figure 4**) and the spindle orientation defects at NEBD that we measured following its depletion (**Figure 3E**), we asked whether LIN-5/NuMA might be required for these forces. We found that most germ cells undergoing prophase after 40 minutes of AID-mediated LIN-5/NuMA depletion showed reduced centrosome movement during prophase and incomplete centrosome separation at NEBD (**Figure 5I-K**). The difference in distance travelled between centrosome pairs was also slightly reduced, although this result was not statistically significant (**Figure 5K**). By contrast, in DHC-1/dynein-depleted cells, net centrosome movement during prophase was similar to control, but centrosome separation at NEBD was more variable (**Figure 5I-K**), with some centrosomes detaching from the nucleus altogether (**Figure 5I and S5A**). These results suggest that LIN-5/NuMA plays a major role in generating the forces that drive and direct centrosome separation in *C. elegans* germ cells, while the dominant role for dynein is to anchor centrosomes to the nuclear surface, where DHC- 1 is also most prominently localized (**Figure S5B;** [56]).

Together, our data support the notion that exclusion of LIN-5/NuMA from the germ cell’s rachis surface, combined with its removal from the cell’s basal cortex in early mitosis biases centrosome movement during prophase to establish spindle orientation parallel to the rachis surface at NEBD.

## Discussion

Here we demonstrate that *C. elegans* germ cells divide parallel to the rachis surface and thus within the plane of the germline tissue during gonad development. These oriented cell divisions are driven by a strong bias in mitotic spindle orientation, with germ cell spindles aligning perpendicular to the cell rachis-basal axis in nearly all cells. This orientation bias is established early in mitosis and relies on the activity of the microtubule motor protein dynein, independently of the cell’s long axis or physical constraints from other tissues in the animal. We find that the dynein regulator LIN-5/NuMA is enriched on all germ cell cortices during interphase, except on their rachis surface, and is redistributed during mitosis, remaining only at lateral cortices along the spindle axis in anaphase. Finally, our results show that germ cell centrioles are positioned near the basal cortex during interphase and that, in prophase, both centrosomes move toward the rachis, positioning themselves at the cell equator and parallel to the rachis surface prior to NEBD and spindle formation.

Our results support a model (**Figure 6**) in which the polarized localization of LIN-5/NuMA throughout the cell cycle establishes germ cell spindle orientation during prophase and maintains it through anaphase. Here, the loss of LIN-5/NuMA from the basal cortex during prophase favors the migration of centrosomes around the nuclear surface, from their starting basal position towards the cell equator, while the lack of LIN-5/NuMA at the rachis surface ensures that neither engages with this cortex, thus ensuring that they reach an orientation parallel to the surface of the rachis at NEBD. The maintenance of LIN-5/NuMA on lateral cortices specifically along the spindle axis after NEBD serves to both stabilize spindle orientation and promote spindle elongation in anaphase. As cells exit mitosis, LIN-5/NuMA returns to the basal cortex, repositioning the disassembling centrosomes basally and preparing cells for the following division. This process, reiterated at each round of cell division, provides a mechanism to ensure spindle orientation parallel to the rachis surface and thus effectively couples germ cell division to planar tissue organization throughout development.

**Figure 6.**
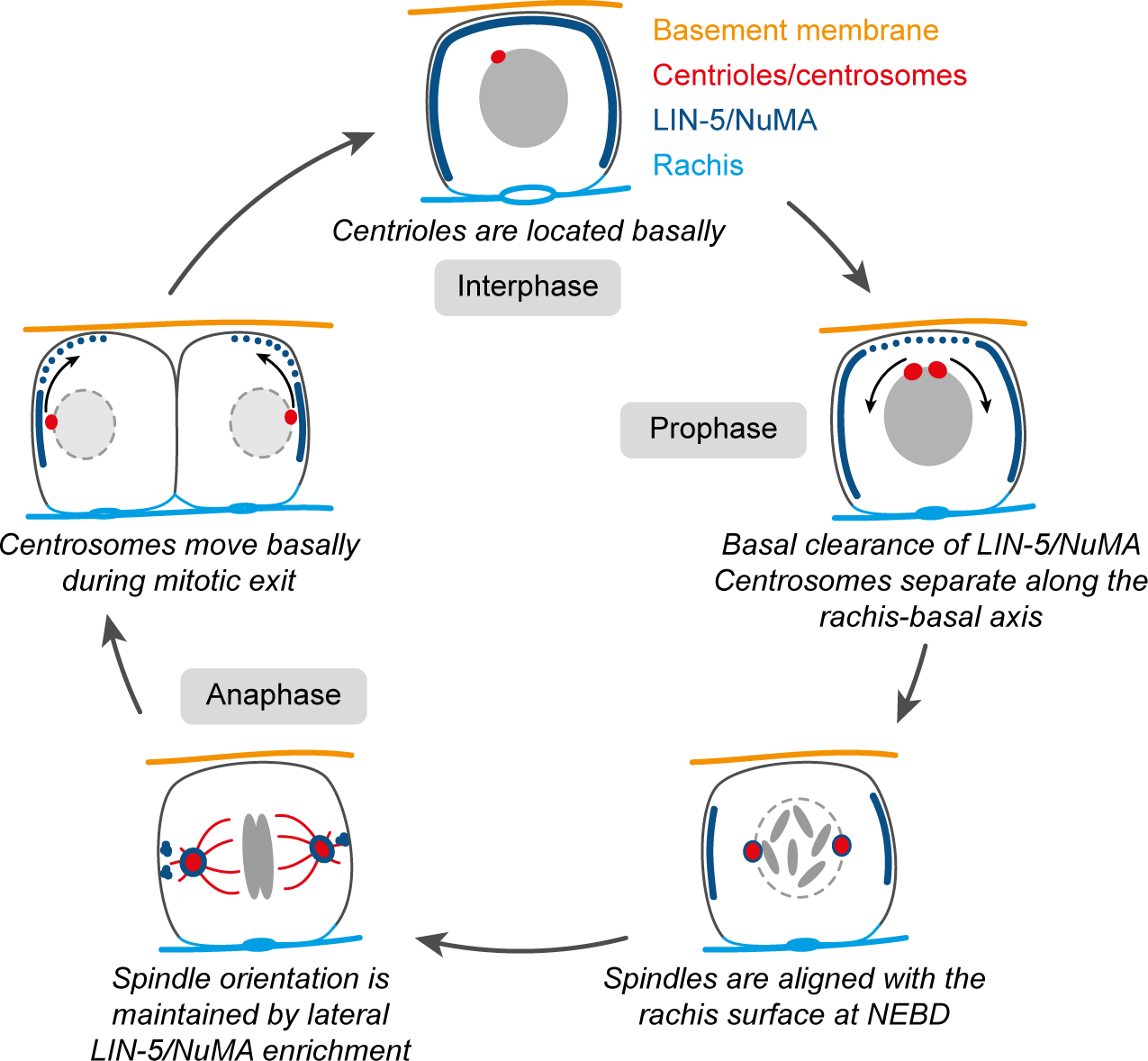
Model for centrosome positioning and spindle orientation during the *C. elegans* germ cell cycle. See main text for details.

This model supposes a key role for centriole positioning in interphase and centrosome dynamics in prophase in establishing proper spindle orientation. In proliferating cells, equal centrosome movement in prophase, from the site of centriole duplication towards opposite sides of the nucleus by NEBD, will typically set spindles up perpendicular to the preceding axis of cell division [57]. Several mechanisms have been described by which cells may overcome this geometric constraint, including tethering one centrosome while the other performs the entirety of migration around the nucleus [58–60], and directing the rotation of centrosome pairs, with the nucleus they flank, after centrosome separation [61]. *C. elegans* germ cells have developed a seemingly different solution to this problem: by shifting the location of the centriole inherited from the preceding cell division basally, equal centrosome migration during prophase can favour a spindle orientation bias parallel to the rachis surface across subsequent rounds of cell division.

While we refer to the ƴ-tubulin puncta we observe in interphase germ cells as centrioles, we do not exclude the possibility that they retain some centrosome and/or microtubule organizing activities. However, we do not detect such foci in interphase germ cells when tracking FP-tagged β-tubulin, and microtubule arrays tend to be non-centrosomal in *C. elegans* meiotic germ cells [62, 63], suggesting that *C. elegans* centrioles are not a major source of microtubule organization outside of mitosis. This then raises the question of how germ cell centrioles come to be basally located in interphase. Our timelapse imaging suggests the interesting possibility that cortical rearrangements during mitotic exit could drag disassembling centrosomes and their associated centrioles basally. High-resolution timelapse imaging to correlate centriole dynamics with both cortical and nuclear movements during mitotic exit are needed to further address this idea.

We note that interphase centrioles are not perfectly aligned with the basal pole of the nucleus, nor is centrosome migration during prophase perfectly symmetric. We infer that ensuring proper spindle orientation at NEBD also requires asymmetries in the forces driving prophase centrosome migration, or that these asymmetries are eventually corrected after NEBD, when spindles undergo rotational movements. In addition, we find that simply setting spindles up parallel to the rachis surface at NEBD is unlikely to ensure that a strong orientation bias persists through anaphase, implying that an active mechanism must also keep spindles within the plane of the tissue throughout mitosis. Our results suggest that the forces dictating prophase centrosome migration and the final orientation of the spindle in anaphase hinge on the spatiotemporal regulation of LIN- 5/NuMA.

How might this regulation then be achieved? Our measurements show that LIN-5/NuMA remains at the germ cell cortex throughout the cell cycle, unlike in most vertebrate cells where its association with the cortex is restricted to meta/anaphase [64]. Its redistribution during prophase suggests that its cortical association is cell cycle dependent. Previous work in the early *C. elegans* embryo and human cell lines demonstrated that LIN-5 and NuMA can be phosphorylated by mitotic kinases, including Cdk1 [65–67], Plk1 [68] and Aurora A [69, 70], any of which could impact LIN-5/NuMA cortical loading in *C. elegans* germ cells. However, loss of these kinases in *C. elegans* leads to pleiotropic defects, and their role in regulating the cortical distribution of LIN- 5/NuMA during germ cell mitosis remains unknown.

LIN-5/NuMA is excluded from the rachis surface in interphase and during mitosis suggesting that its cortical distribution is also influenced by germ cell rachis-basal polarity. A defining feature of the *C. elegans* germline architecture is the presence of actomyosin-rich intercellular bridges connecting germ cells to the rachis on their apical side [29, 33]. While it is formally possible that the presence of a rachis bridge, and the gap it generates in the germ cell rachis membrane, limits the association between LIN-5/NuMA and this cell surface, several reasons suggest that this is not a major determinant of LIN-5/NuMA cortical polarization. First, rachis bridges are effectively closed during mitosis, yet association between LIN-5/NuMA and the rachis surface does not change (**Figure 4D-E**). Furthermore, LIN-5/NuMA is excluded from the entirety of the rachis surface, including regions well away from the rachis bridge (**Figure 4A-C**). Instead, as rachis bridges are dynamic and under tension [31, 32, 39], and anisotropies in actomyosin tension can influence the activity of force generators and spindle orientation in other contexts [71], it is possible that the enrichment of actomyosin contractility regulators and/or the mechanical properties of the rachis cortex locally restrict the amount of LIN-5/NuMA.

Finally, during mitosis, LIN-5/NuMA is enriched on lateral germ cell cortices, but only those behind the spindle poles, where it appears as transient foci. This localization pattern is similar to that seen in several other cell types where clustering of the force generating machinery has been implicated in spindle orientation [3, 72, 73]. In addition, evidence from several systems including the *C. elegans* embryo [74], budding yeast [75], *Drosophila* [76] and mammalian cells [77, 78] suggests that, in some cases, astral microtubules direct the cortical deposition of force-generating complexes to focus pulling forces at sites of microtubule-membrane contact. It is therefore possible that maintenance of spindle orientation during germ cell mitosis uses a similar mechanism. Under this scenario, exclusion of LIN-5/NuMA from the germ cell rachis surface should be sufficient to maintain spindles parallel to this surface, while spindle-driven focusing of the force generating machinery gives spindles flexibility within the plane of the tissue to respond to other cues which could contribute to spindle orientation along the tissue D/P axis. Future work is necessary to determine the nature of these cues and the precise mechanism by which force generation is consolidated on germ cell lateral membranes.

The organization of epithelial tissues typically depends on cell-cell junction complexes, which polarize cells along their apical-basal axis and often coordinate spindle orientation during mitosis to enable planar tissue expansion. The notable absence of cell-cell junctions between *C. elegans* germ cells [29, 30] hinted that the maintenance of tissue organization during development relies on a different mechanism. Our work suggests that *C. elegans* germ cells undergo oriented cell divisions by localizing their centrioles basally in interphase and then controlling the distribution of cortical force generators in both time and space to ensure that the mitotic spindle is established and then maintained within the tissue plane. This work presents an alternative model by which spindle orientation and oriented cell division can be achieved and highlights the mechanistic plasticity of these processes during animal development.

## Materials and Methods

### *C. elegans* strain maintenance

*C. elegans* animals were maintained at 20°C on nematode growth medium (NGM) and fed with *Escherichia coli* strain OP50 according to standard protocols [79]. For all experiments, late L4 stage animals were identified by size and vulva morphology and individually collected from these plates except for the dataset of Figure 1, where different developmental stages were obtained by synchronizing larvae at the L1 stage. Synchronized L1 larvae were obtained by sodium hypochlorite treatment [80]. Briefly, gravid hermaphrodites were dissolved in a solution of 1.2% sodium hypochlorite and 250 mM sodium hydroxide. Pelleted embryos were washed 3 times in M9 buffer (22.04 mM KH_2_PO_4_, 42.27 mM Na_2_HPO_4_, 85.55 mM NaCl, 1 mM MgSO_4_), and allowed to hatch for 24 h at 15°C in M9 buffer. Animals at the L3, late L4 and adult day 1 stages were obtained by inoculation of synchronized L1 larvae on NGM plates containing 1 mM isopropyl β-d-thiogalactoside and 25 μg/ml carbenicillin and fed for respectively 26-30, 40-48 and 72 hours with *E. coli* strain HT115 transformed with the empty RNA interference (RNAi) vector L4440. The strains used in this study are listed in Table S1.

### Generation of mAID::mNG-tagged LIN-5

The plasmid used for tagging *lin-5* with mNeonGreen (mNG) and mAID was constructed as follows. Homology arms in the coding sequence of *lin-5* and in the 3’-UTR region were amplified using genomic DNA as a template. mAID was amplified from the plasmid pMK290 ([81]; Addgene: #72828). mNG was amplified from a plasmid containing *mNG-mom-5* [82]. The self-excising cassette (SEC) was amplified from the plasmid pDD287 ([83], Addgene #70685). The plasmid backbone was amplified from the pDD287 plasmid. These fragments were assembled using NEBuilder to construct the donor plasmid, pTN26 (*lin-5::mAID::mNG*). For Cas9 and sgRNA expression, a guide RNA sequence (5′-GTCCAAGAAAAAGAACCGTC-3′) targeting the C- terminal coding region of the *lin-5* gene was used, as in [82]. The plasmid, pTN27 (sg *lin-5*), was constructed by inverse PCR with the plasmid pDD162 ([84], Addgene #47549). Editing of the *lin- 5* locus was performed as described previously [85]. Briefly, 25 ng/µl of pTN26 and 5 ng/µl of pTN27 were injected into the gonad of N2 animals with the control injection markers pCFJ90 (*Pmyo-2::mCherry*, Addgene #8984,[86]) and pCFJ104 (*Pmyo-3::mCherry*, Addgene #19328 [86]). The site of mAID::mNG insertion was verified by PCR on the genomic DNA of homozygous progeny. All oligonucleotide sequences are reported in Table S2.

### Worm mounting and live imaging

Animals were anesthetized in M9 buffer containing 0.04% tetramisole (Sigma, Cat #L9756) and transferred to a 3% agarose pad, molded with *C. elegans* stage matching-sized grooves made by a custom microfabricated silica plate, as described previously [37, 38, 52]. A glass coverslip was placed onto the pad, the chamber was backfilled with M9 buffer containing 0.04% tetramisole and sealed using VaLaP (1:1:1 Vaseline, Lanolin, and Paraffin). Images were acquired at room temperature (∼20°C) on either a Zeiss Cell Observer spinning disk confocal microscope (Zeiss inverted Cell Observer with a Yokogawa CSU-X1 confocal scanner, controlled by Zen software, using a Zeiss 63x/1.4 NA Plan Apochromat DIC (UV) VIS-IR oil immersion objective, 488 nm (30mW) and 561 nm (50mW) solid-state lasers, with a quad pass 466/523/600/677 emission filter and a Zeiss AxioCam 506 Mono camera) or a Nikon CSU-X1 spinning disk confocal microscope (Nikon TI2-E inverted microscope with a Yokogawa CSU-X1 confocal scanner, controlled by NIS-Elements software, using either a Nikon Plan Apo Lambda 60x/1.4 NA Oil immersion objective or a Nikon Apo 40x/1.25 NA water immersion objective, 488 nm (100mW) and 561 nm (100mW) solid-state lasers, with a dual band pass Chroma 59004m filter or single pass filters EM525/50, EM605/55, EM700/75, and a Hamamatsu ORCA-Fusion BT sCMOS camera). Detailed imaging conditions are reported in Table S3.

### Auxin treatment

A 400 mM Auxin stock in ethanol was prepared from natural auxin indole-3-acetic acid (IAA, Sigma Cat #I3750) as described in [51]. Auxin was added at final concentration of 1mM to the NGM solution before pouring plates. Plates were inoculated with OP50 and left at room temperature in the dark for 2 days before use. L4 stage animals were collected from OP50 seeded NGM plates, washed in 100 μL of M9 buffer and then transferred to OP50 seeded NGM-Auxin plates for either 40 minutes or 6 hours at room temperature in the dark prior to mounting and imaging. The level of depletion was quantified by measuring the mean fluorescence intensity of DHC-1::AID::GFP and LIN-5::mAID::mNG in control and AID-treated animals in Fiji 1.52v [87]. Three ∼200 µm^2^ rectangles were drawn manually across the mitotic region of the gonad and the average intensity was measured in 3 z-slices around the middle of the gonad. Measurements are reported as the mean value per gonad after autofluorescence was subtracted. Autofluorescence was measured using the same approach but on gonads from animals carrying only mCH::β- tubulin (strain ARG3).

### Gonad explants

Three L4 stage larvae, raised on OP50-seeded NGM plates, were transferred into a ∼5 µl drop of meiosis medium (0.5 mg/ml Inulin (Sigma, Cat #I2255), 25 mM HEPES pH 7.5 (Wisent, Cat #600- 032-CG), 60% Leibovitz’s L-15 Media (Wisent, Cat #323-050-CL), 20% FBS heat inactivated (Wisent, Cat #090-150, lot 112740)) [40], on a glass slide patterned with 14mm x 14mm wells (Fisher Scientific, 30-2066A-BROWN 3 SQUARE 14mm with Bars Epoxy autoclavable). Using 25-gauge needles (BD Precision Glide #CABD305122), animals were cut below the pharynx, extruding at least one gonad arm into the medium. A coverslip was then gently placed over the drop and sealed with VaLaP. With this method, ∼1/3 of explants remained intact, as assessed by germ line morphology, and were kept for time-lapse imaging.

### Germ cell centrosome tracking and scoring of mitotic events

Image registration, centrosome tracking and pairing, and scoring of mitotic events (e.g. nuclear envelop breakdown (NEBD)) were performed using CentTracker, as described previously [37, 38]. Briefly, z-stack images were registered to correct for sample movement over time. Centrosomes were tracked in Fiji 1.52v [87] using the plug-in TrackMate v6.0.0 [88] and their x-y-z-t coordinates were processed using a trainable, machine-learning–based approach to retain true pairs of centrosomes (captures ∼70% of centrosome pairs per germ line in wild-type conditions). For experiments other than those in Figure 1, the true pair processing step was done manually to retain the maximum number of centrosome pairs possible (close to 100% per germ line). The centrosome-to-centrosome distance (spindle length) overtime was then used to define four mitotic events: the start of centrosome separation in prophase, NEBD, prometaphase-metaphase, and anaphase onset, as described [37, 38, 52].

### Cell surface rendering and cell long axis extraction

To determine germ cell shape, volume, and orientation in 3D, germ cell membranes, marked by the fluorescent protein (FP)-tagged PH domain of rat PLC1∂ [46], were rendered using the Cells tool in Imaris (version 9.2.1, Oxford Instruments) [89], with the following parameters: cell detection type = cell membrane, smallest diameter = 3 µm, cell membrane detail = 0.324 µm and cell filter type = local contrast. Intensity and quality filters were applied manually, on a per gonad basis, to be as close as possible to the cell membrane marker (as visualized using the Surpass viewer in Imaris). Cell statistics (cell volume, cell position, cell ellipsoid axis orientations, cell ellipsoid axis lengths, and cell ellipticity indices) were computed in Imaris and data were exported as CSV files, which were then imported into Julia (version 1.10.4) [90]. Using the centrosome tracking data (as described above), the x-y-z-t coordinates of the spindle midpoint were used to extract the corresponding cell surface renderings from the Imaris output file. All remaining cells were considered interphase cells. Interphase cells were also filtered to select tracks of at least 10 timepoints, with a mean volume greater than the smallest newly divided daughter cells and lesser than the largest mitotic cells (75 µm^3^ < cell volume < 250 µm^3^), and where cell volume was relatively stable over the 10-frame track duration (standard deviation < 20 µm^3^ and mean rate of change in volume over time within ± 2 µm^3^/min). To select cells with a stable long axis, the orientation of the ellipsoid major axis at each timepoint was used to calculate the mean orientation of the long axis over time, and cells were excluded when frame-to-frame orientations deviated significantly from this mean (mean difference between measured orientations and the calculated mean orientation > 20°) or when the standard deviation of long axis orientation exceeded 20°. For selected interphase cells and mitotic cells, cell shape was described by the ellipticity prolate index (*e_prolate_*) of the fitted ellipsoid:

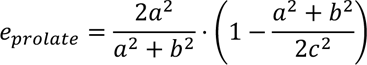

where *a*, *b* and *c* are the lengths of the three ellipsoid axes. For mitotic cells, the orientation of the cell long axis was defined as the mean orientation of the longest axis during 5 timepoints prior to anaphase onset (AO -4 to AO); for interphase cells, the orientation of the cell long axis was defined as the mean orientation during the first 10 tracked timepoints.

### Defining the gonadal D/P axis

The D/P axis was defined as previously described [37]. Briefly, a rectangle encompassing the distal region of the gonad arm at the center z-slice was drawn. The D/P axis was defined as the vector joining the midpoints of the short sides of the rectangle. In explants, the gonad was not necessarily oriented parallel to the image plane and the rectangle used to define the D/P axis was determined locally along a 30 µm stretch of the gonad surrounding each dividing cell, and adjusted as needed to accommodate gonads that were tilted relative to the imaging plane.

### Rachis and gonad surface rendering

To render the rachis surface, z-stack timelapse images were imported into Imaris and the rachis was delineated using the surface rendering tool. A gaussian filter was applied with surface details set to 0.75 µm and the rendering threshold was adjusted manually, on a per gonad basis, to be as close as possible to the rachis surface (as visualized using the Surpass viewer in Imaris). A filter was then applied to keep only the largest object rendered per time point. The rendered surface was exported to Fiji as a binary hyperstack TIF and a custom macro was used to fill, in 3D, holes in the rendered surface. The processed TIF was then reimported into Imaris, and the rachis surface was re-rendered using the same parameters as before. The final rendered surface was exported as series of vertices and edges, forming a meshwork of triangles, in Virtual Reality Modeling Language format (WRL), which was converted to a CSV file using a custom Python 3 script [91] for use with MATLAB (version 2020b) [92] or imported directly to Julia. The gonad surface was rendered using the same method, except that surface details were set to 2 µm and the hole filling step was not necessary. Gonad surface renderings were generated using the FP-tagged PH domain or β-tubulin signal, both of which are expressed solely in the germ line.

### Determining the orientation of the rachis or gonad surface for each cell

Rachis or gonad surface renderings were imported into MATLAB or Julia, where they were paired with centrosome tracking and/or cell surface rendering data. A patch of rachis or gonad surface meshwork triangles was extracted from the rendered surface for each cell at each timepoint, by calculating the Euclidean distance between the spindle midpoint and/or fitted ellipsoid centroid and the centroid of all surface triangles. Triangles were then sorted by distance and the sum of the area of all triangles within 0.01 µm of the spindle midpoint or ellipsoid centroid was calculated. This process was performed iteratively, increasing the distance from the spindle midpoint or ellipsoid centroid by 0.01 µm until the cumulative measured area reached at least 50 µm^2^ or the distance reached 8 µm. If a 50 µm^2^ net area was not reached within 8 µm, no patch was defined for that timepoint. A clustering analysis of the selected triangles’ centroids was performed to exclude timepoints where non-contiguous patches were identified. The normal vector of the obtained patch (the rachis or gonad surface normal) was defined as the sum of the normal vectors for all triangles within the patch.

### Measuring spindle and cell long axis orientations

The orientation of the spindle or cell long axis relative to the rachis surface was calculated as the angle between the rachis normal and the spindle vector or cell long axis in 3D as follows:

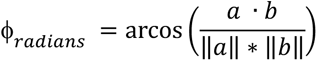

where a and b are the 2 vectors in 3D. φ was converted from radians to degrees. Angles greater than 90° were normalized to a 0° to 90° range by taking the corresponding acute angle ( φ = 180 - φ ). The same calculation was performed when determining the orientation of the spindle or cell long axis to the D/P axis, or the orientation of the spindle to the gonad surface.

Spindle orientation measurements were compared to a theoretical random distribution, calculated as described previously [37, 93]. Briefly, the probability of a vector occupying a given angle around its midpoint, is proportional to the surface area of a sphere, centered on the vector midpoint, at that angle. For example, a sphere with a radius of 1, has circumference C at angle φ from its polar axis, as given by:

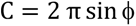

Therefore, the surface area, Aφ, between the angle φ and the polar axis (0°) and its symmetrical counterpart -φ (with 0° ≤ φ ≤90°) can be calculated as:

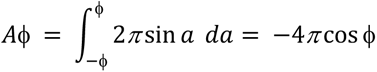

The fraction fφ of the total cortical surface area can then be calculated as:

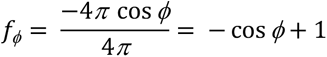

The rachis normal, D/P axis or the cell long axis, were considered as the sphere’s polar axis, depending on the spindle orientation being assessed. Deviation from the theoretical random distribution was assessed as described [93], with λ_p_ and λ_o_ indicating how enriched spindle orientations are parallel or orthogonal to the reference axis, respectively.

To calculate the theoretical range of spindle orientations that would preclude spindle-rachis contact in anaphase, the average distance between the spindle midpoint and the rachis surface (here modelled as the best-fit plane to the rachis surface patch) and the average spindle half-length at the end of anaphase spindle elongation were considered as two sides of a right triangle (short side and hypotenuse, respectively) and used to calculate the angle between the spindle vector and rachis normal assuming that a line on the rachis plane formed the third side of the triangle.

To predict the range of spindle orientations to the rachis normal at anaphase onset if the only constraint were spindle orientation at NEBD, the measured angles between the spindle vector and the rachis normal from NEBD to anaphase onset for the cells analyzed in Figure 1 were used to calculate the frame-to-frame change in spindle angle, with the direction of rotation (towards or away from the rachis normal) preserved (hereafter rotation). We note that because spindle angles were reported as within 0° to 90° of the rachis normal, the range of rotations used here underestimates the true extent of spindle movement. A basic simulation was then run in MATLAB in which a starting angle and a duration of mitosis (NEBD to anaphase onset) were drawn at random from all measured values, and a set of rotations, with the number determined by the duration of mitosis (one rotation per frame), were drawn at random from all measured rotations for all cells. Net rotation during mitosis was calculated by summing the set of randomly drawn rotations and the resulting angle at anaphase onset was calculated as the angle at NEBD + net rotation, with obtuse anaphase angles returned to within 0° to 90° of the rachis normal, as above. The simulation was run 1000 times, and the output was used to plot the cumulative distribution of predicted anaphase angles relative to the observed distribution and the distribution of predicted net rotations versus measured net rotations.

### Cortical fluorescence intensity measurements

To measure cortical fluorescence, a series of vectors were generated using the spindle midpoint (for mitotic cells) or the cell center (defined manually in Fiji by placing a point ROI at the center point of the nucleus, as identified by the exclusion of cytoplasmic fluorescence) for interphase cells. Interphase cells were selected by size (≳125 µm^3^) and the absence of neighboring mitotic cells. For mitotic cells, cortical fluorescence was measured along the spindle vector, the gonad surface normal (generated from the gonad surface rendering, as described above) and a third vector orthogonal to both. For interphase cells, cortical fluorescence was measured along the gonad surface normal, a vector parallel to the gonadal D/P axis passing through the cell center and a third vector orthogonal to both. A series of flattened ellipsoids (2 µm high x 2 µm wide x 0.5 µm deep) were propagated along each vector, every 0.25 µm, for a total distance of 6 µm on either side of the spindle/cell midpoint. The orientation and position of each ellipsoid was calculated using a custom MATLAB script, imported into Fiji, and a custom macro was used to draw the corresponding 2D ROI where the ellipsoids intersected with each z-slice. The area and raw integrated density (RID) for each 2D ROI for each ellipsoid was summed and converted to a mean fluorescence intensity (FI; total RID/total area) per ellipsoid. A custom MATLAB script was then used to plot FI, relative to distance along each vector, for both the FP-tagged PH membrane marker and LIN-5::mNG. The position of the peak FI for the membrane marker was used to identify where the vector intersected with the cell cortex and the corresponding LIN-5::mNG FI value was extracted. Background (the average minimum FI along the cell orthogonal vector) was subtracted and LIN-5::mNG FI was normalized to its respective membrane signal. For mitotic cells, LIN-5::mNG measurements were made at anaphase onset (determined by centrosome tracking, as described above) and corrected for photobleaching, according to when, relative to the start of imaging, each cell entered anaphase. Briefly, a bi-exponential function was fit to the change in FI over time in a rectangle (∼20 µm x 100 µm) drawn on a maximum intensity projection through the gonad that encompassed the mitotic zone. The reverse of this function was then applied to the measured data. For interphase cells, FI measurements were made at the first frame of the movie. For mitotic cells, the reported spindle and orthogonal vector values show the mean FI for both cortices along each vector. To avoid measuring centrosome-localized LIN-5::mNG, timepoints were excluded if the distance between the centrosome and cortex was less than 1.5 µm. We also excluded cells if centrosome-localized LIN-5::mNG in neighboring mitotic cells was captured by the line scan. For interphase cells, lateral measurements reflect the mean of all 4 cortices (2 D/P + 2 orthogonal cortices).

### Analysis of centrosome movement and position in prophase

Centrosome tracks were generated using TrackMate, as described above, and the x-y-z coordinates for each centrosome were expressed relative to the centroid of the nucleus. The nucleus centroid was determined either by tracking nuclei in TrackMate using the histone maker or manually in TrackMate using the exclusion of cytoplasmic fluorescence (FP- and AID-tagged LIN-5 and DHC-1) to identify the nucleus. Frame-to-frame centrosome movement was assessed by assuming that centrosomes move around the surface of the nucleus and calculating the arc length as follows: d = φ * r, where φ = the central angle between centrosome positions at adjacent timepoints and r = nuclear radius (as defined by the distance between the centrosome and the center of the nucleus).

### Measuring centriole position in interphase cells

The x-y-z coordinates of the centroid of select nuclei were used to generate a 10 µm-thick stack, oriented orthogonally to the gonadal D/P axis, and centered on each nucleus, using the “Reslice” tool in Fiji. Maximum intensity projections of the resliced stack were used to create a cross-sectional view of the gonad and a 3 µm-thick line was drawn from the center of the gonad, through the center of the nucleus towards the basal surface of the gonad, approximating the gonadal radial axis. Fluorescence intensities for both FP-tagged histone and ƴ-tubulin were plotted and used to define the diameter of the nucleus and the position of the centrioles on the nucleus, along the gonadal radial axis. Centriole position was expressed as a percent of nuclear diameter. To exclude mitotic cells, only cells where a single ƴ-tubulin focus was visible were analyzed.

### Statistical analysis and figure assembly

Statistical analysis was performed in MATLAB or Julia. One-sample Kolmogorov-Smirnov tests (MATLAB kstest) with Bonferroni corrections [94], as needed, were used to compare the distribution of measured angles to the theoretical random distribution. Two-sample Kolmogorov– Smirnov tests (MATLAB kstest2) with Bonferroni corrections, as needed, were used to compare the distribution of measured angles between two samples. Kruskal–Wallis tests with a Tukey– Kramer post hoc test (MATLAB kruskalwallis and multcompare) were used to compare multiple independent samples means. Two-tailed Student’s t-tests (MATLAB ttest2) were used to compare two independent samples, except when comparing the duration of mitosis between cells in explants versus *in situ*, where a two-tailed Student’s t-test with unequal variance was used (MATLAB ttest2 with Vartype = unequal). A one-sample t-test (MATLAB ttest) was used to test if centrosome midpoint displacement during prophase (**Figure 5G**) had a mean different from zero. Bartlett multiple-sample tests for equal variances (MATLAB vartestn) were used to compare the variance of centrosome separation in controls and LIN-5 or DHC-1 depleted conditions, followed by a two-sample F-test for equal variances (MATLAB vartest2) to identify which pairs were significantly different (**Figure 5J**). All summary statistics are provided by figure panel in Table S4. Figures were assembled using Adobe Illustrator. Graphs were generated in MATLAB or Julia, saved as PDFs and imported into Illustrator to generate high resolution, vector-based graphics. All representative images were processed in Fiji (scaled to the same brightness and contrast settings, pseudo colored and cropped) and exported as RGB TIFs. Images were re-sized and/or cropped Illustrator to fit the final figure panel.

## Acknowledgments

We are grateful to Drs. Arshad Desai (University of California, San Diego), Amy Maddox (University of North Carolina, Chapel Hill), Pablo Lara-Gonzalez (University of California, Irvine) and Benjamin Lacroix (Université Paris Cité, Paris) for sharing strains and reagents. We also thank Christian Charboneau of IRIC’s Bio-imaging facility and the staff at McGill’s Advanced BioImaging Facility (ABIF) for technical assistance, and members of the Labbé, Gerhold and Weber labs for stimulating discussions and advice. Some strains were provided by the *Caenorhabditis* Genetics Center, which is funded by the NIH Office of Research Infrastructure Programs (P40 OD010440). The mNG-mom-5 plasmid was kindly provided by Drs Jennifer Heppert and Bob Goldstein. RMZ holds doctoral scholarships from the Natural Sciences and Engineering Research Council of Canada (NSERC) and the Fonds de recherche du Québec – Nature et Technologies (FRQ-NT). This work was funded by grants from the Canadian Institutes of Health Research (PJT-153283) to JCL and ARG and JSPS KAKENHI (23K05836) to TN.

## Author contributions

Conceptualization, R.M.Z, V.P., J-C.L., A.R.G; methodology, R.M.Z, V.P., J-C.L., A.R.G; software, R.M.Z, V.P., A.R.G; formal analysis, R.M.Z, V.P., T.N.; investigation, R.M.Z, V.P., T.N.; data curation, R.M.Z, V.P.; writing – original draft, R.M.Z, J-C.L., A.R.G; writing – review & editing, R.M.Z, V.P., T.N., J-C.L., A.R.G; visualization, R.M.Z, V.P., A.R.G; funding acquisition, J-C.L., A.R.G.

## Supplemental information titles and legends

Document S1. Figures S1-S5 and legends. Tables S1 – S3. Supplemental video legends.

Table S4. Excel file containing summary statistics for all figures.

Video S1. Centrosomes and rachis bridge dynamics during germ cell mitosis. Related to Figure 1.

Video S2. LIN-5::mNG at the germ cell basal cortex during prophase. Related to Figure 4.

Video S3. LIN-5::mNG foci on the lateral cortex of a germ cell prior to anaphase. Related to Figure 4.

Video S4. Centrosomal LIN-5::mNG is redistributed to the basolateral cortices at the end of mitosis. Related to Figure 4.

---

**Figure S1.**
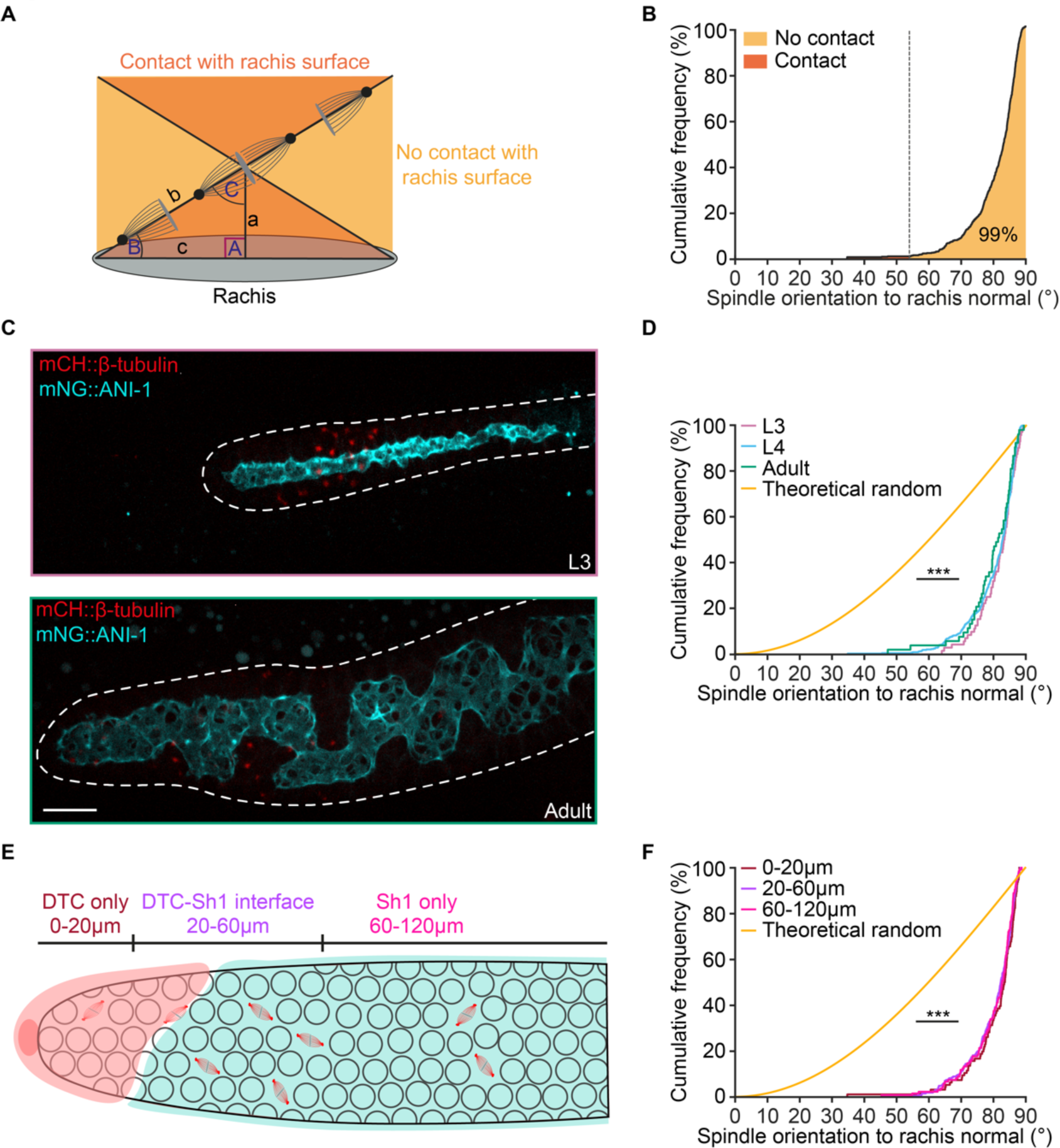
Germ cell spindles orient parallel to the rachis surface irrespective of developmental stage and distance to the niche. (A) Schematic representation of the calculation used to predict the angles to the rachis normal that would result in one of the two spindle poles making contact with the rachis surface (grey) in anaphase. a = mean distance between the spindle midpoint and rachis surface. b = mean spindle half-length at the end of anaphase elongation. C = angle between spindle vector and rachis normal formed by the right triangle with side a and hypotenuse b. (B) Cumulative distribution of spindle angles to the rachis normal measured during anaphase in L4 animals. The vertical dotted line marks 53°, beyond which 99.6% (n = 503) of mitotic spindles are found. (C) Maximum intensity projection of a distal gonad arm in an L3 larva (top) and an adult animal (bottom) expressing mCH::β-tubulin (red) and mNG::ANI-1 (cyan). Scale bar = 10 µm. (D) Cumulative distribution of spindle angles to the rachis normal measured during anaphase in L3, L4 and adult animals. L4 data are reproduced from (B). (E) Schematic representation of the distal gonad arm of an L4 animal depicting the approximate position of the DTC niche-only (0–20 µm), DTC niche-Sh1 sheath cell interface (20–60 µm) and Sh1 sheet cell-only (60–120 µm) regions of the gonad, as measured from the distal-most tip of the tissue [1–3]. (F) Cumulative distribution of spindle angles to the rachis normal from data in (B), divided by gonad region as defined in (E).

**Figure S2.**
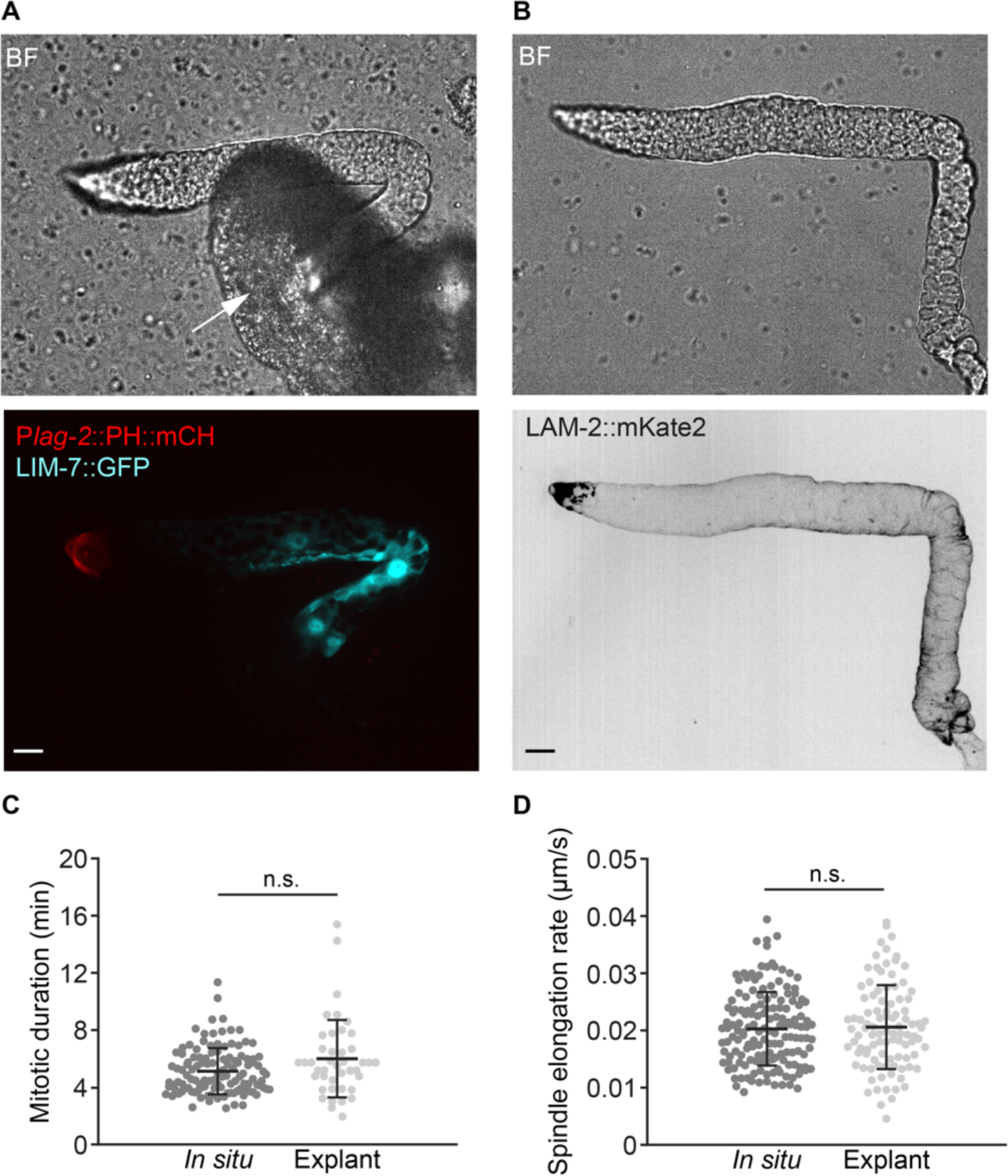
*C. elegans* gonad explants retain somatic gonadal cells and proper mitotic spindle dynamics. (A-B) Bright field images (top) and maximum intensity projections (bottom) of gonad arm explants from L4 animals expressing markers of the DTC (P*lag-2*::PH::mCH, red) and sheath cells (LIM-7::GFP, cyan; A) or the basement membrane (LAM-2::mKate2; B). The white arrow in (A) points to a fragment of the extruded gut. Scale bars = 10 µm. (C-D) Comparative measurements of the duration of germ cell mitosis (C) and spindle elongation rates during anaphase (D) made *in situ* and in gonad explants. Bars indicate average ± standard deviation.

**Figure S3.**
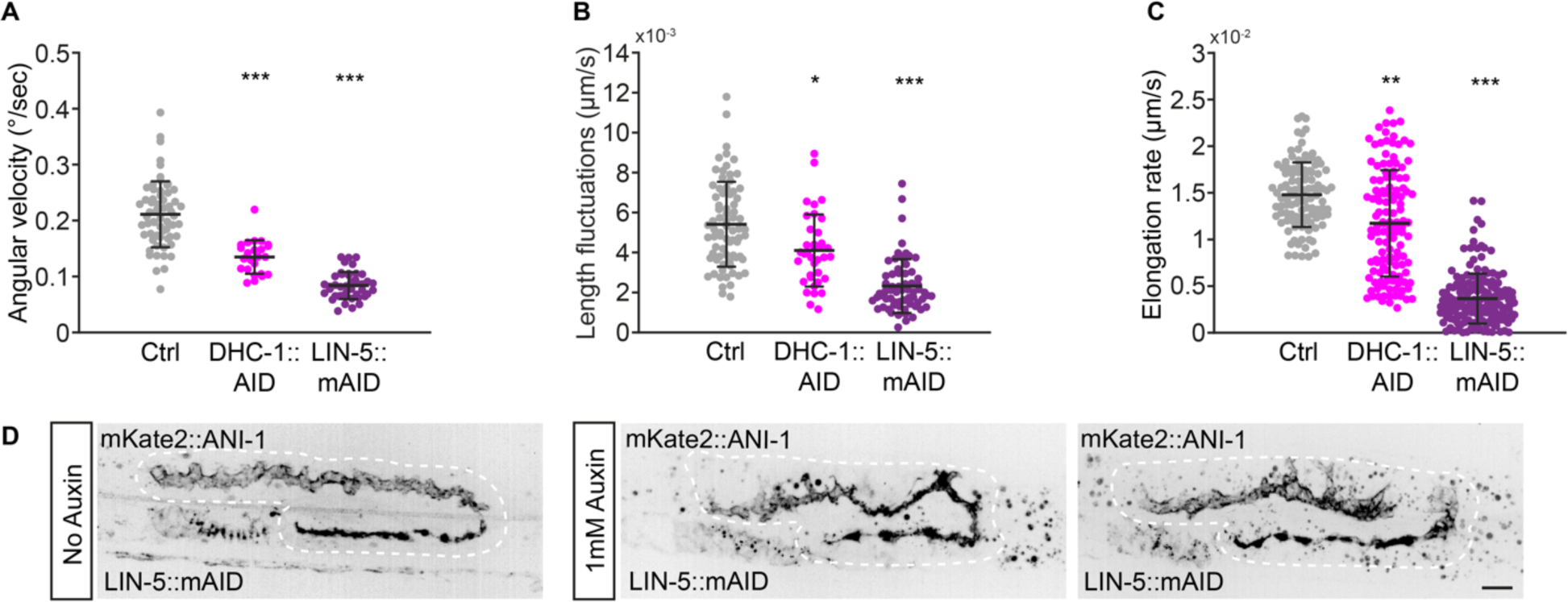
LIN-5/NuMA and DHC-1/dynein are required for proper spindle dynamics and rachis organization. (A-C) Measurements of mitotic spindle angular velocity (A) and spindle length fluctuations (B) in prometa/metaphase, and rate of spindle elongation during anaphase (C) in germ cells from control animals and animals depleted of DHC-1/dynein or LIN-5/NuMA in the germ line by a 40-minute auxin treatment. (D) Maximum intensity projections of gonad arms from L4 larvae expressing endogenously tagged mKate2::ANI-1 to mark the rachis, without (left) and after (right) a 6-hour auxin treatment to deplete LIN-5::mAID. Scale bar = 10 μm.

**Figure S4.**
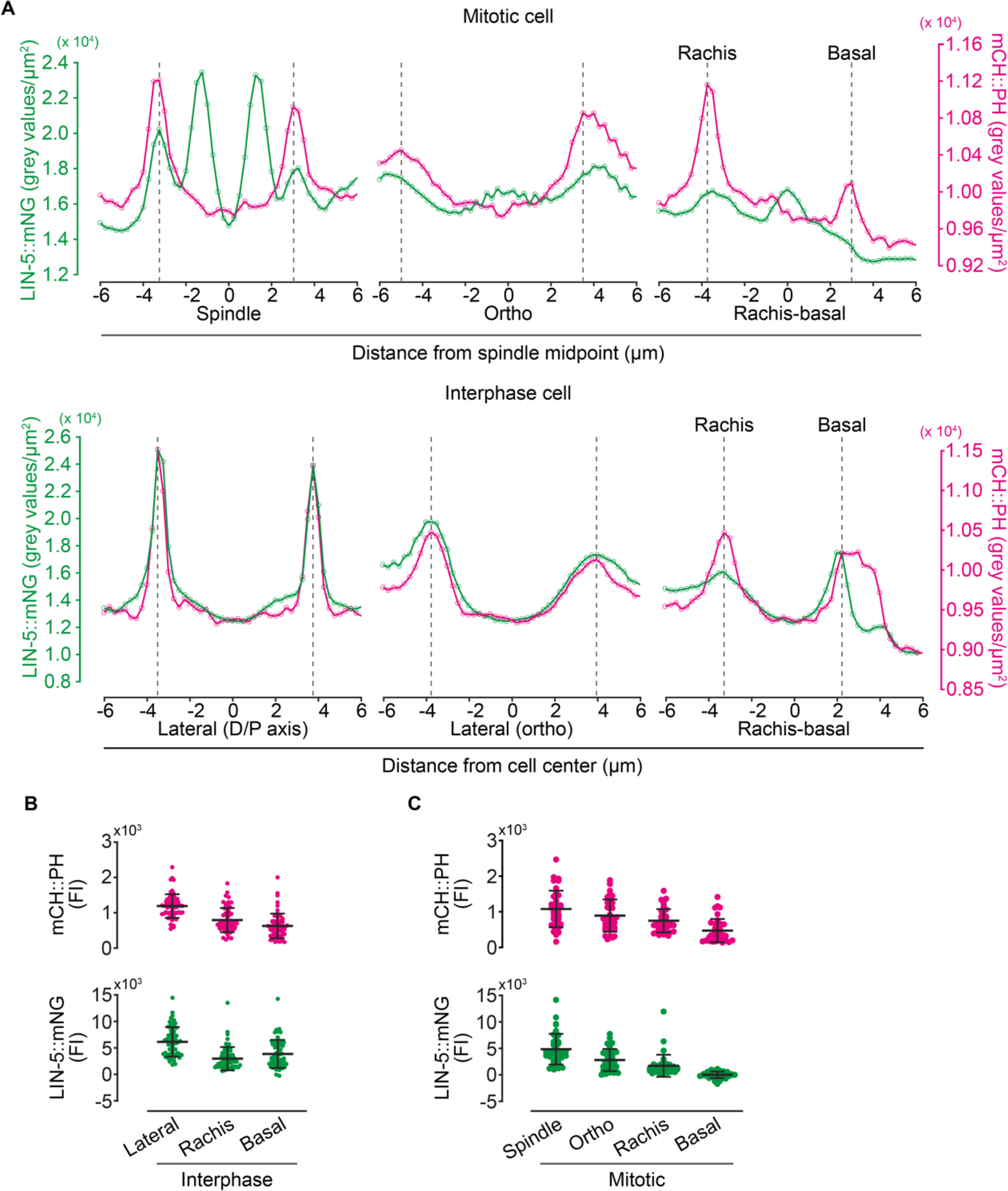
Measuring LIN-5/NuMA cortical localization in germ cells. (A) Representative line scans showing the raw fluorescence intensity (FI) measurements of LIN-5::mNG (green, NuMA) and mCH::PH (magenta, membrane) along the different axes indicated (spindle, ortho and rachis- basal for the mitotic cell (top) and lateral (D/P), lateral (ortho) and rachis-basal for the interphase cell (bottom)). The peak in membrane FI used to extract the corresponding LIN-5::mNG FI is shown for each cortex. (B-C) FI measurements of mCH::PH (top) and LIN-5::mNG (bottom), after background subtraction and averaging across cortices along the same vector (all lateral cortices for interphase cells (B) and spindle and ortho cortices for mitotic cells (C)).

**Figure S5.**
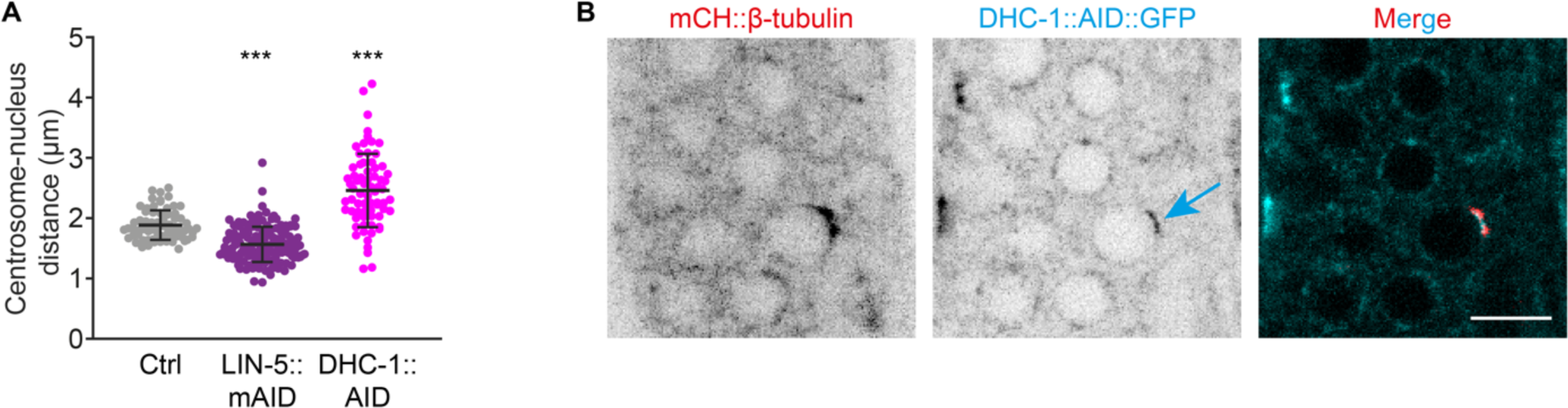
Dynein favours centrosome attachment to the nucleus. (A) Measurements of centrosome-to-nucleus distance in germ cells of L4 animals subjected or not to a 40-minute auxin treatment to deplete LIN-5/NuMA or DHC-1/dynein. Centrosomes tend to closer to the nuclear center following LIN-5/NuMA depletion and further from the nuclear center following DHC- 1/dynein depletion. (B) Maximum intensity projections of mCH::β-tubulin (red) and DHC- 1::AID::GFP (cyan) in the distal gonad of an L4 larvae, showing the relative enrichment of DHC- 1/dynein around nuclei at its co-localization with centrosomes early in prophase (arrow). Scale bar = 5 μm.

**Table S1.**
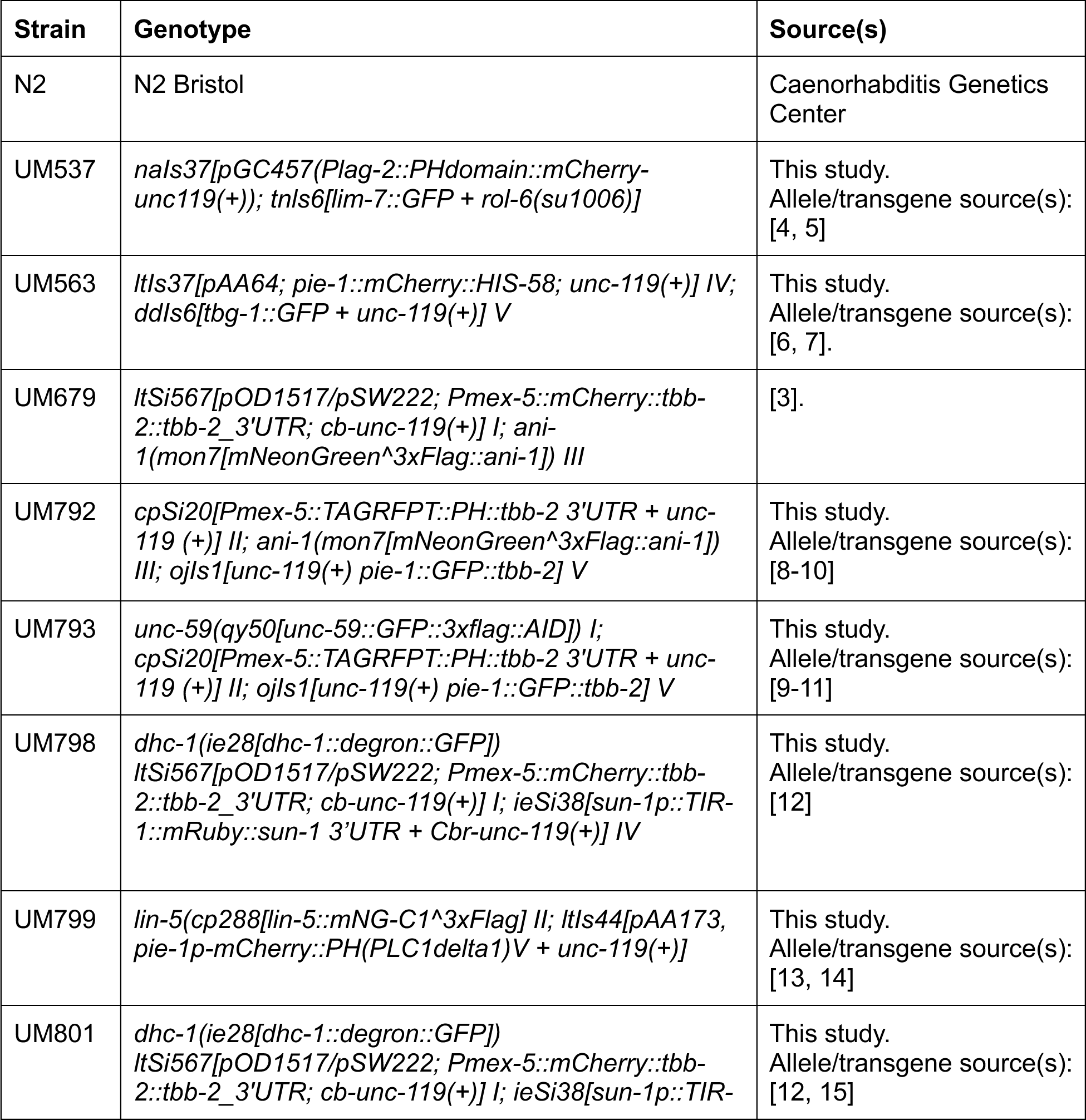
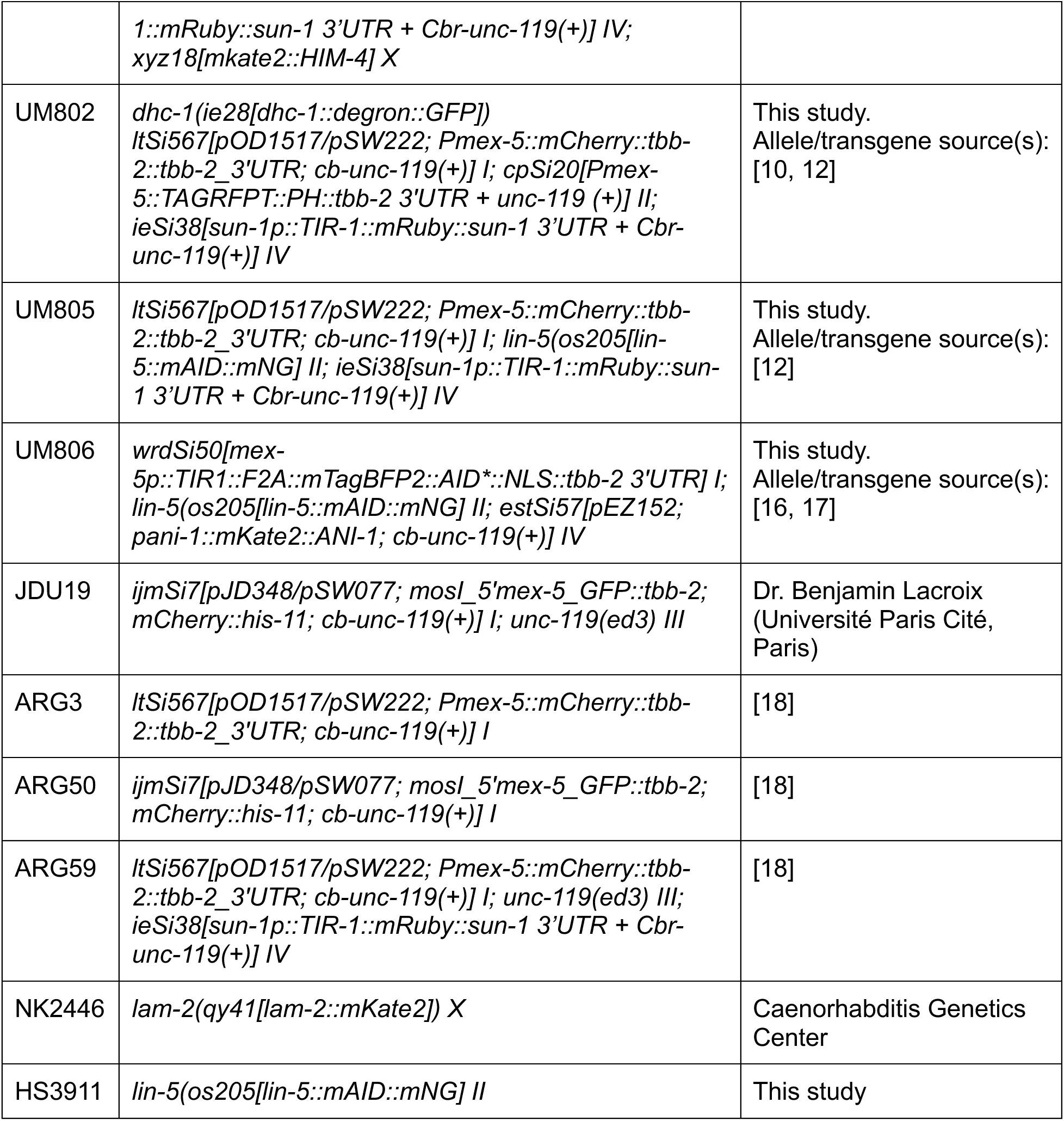
List of C. elegans strains used in this study. Unless otherwise noted, all strains were constructed as part of this study. References indicate the source(s) for alleles and/or transgenes used in strain construction. During strain construction, *unc-119(ed3)* was not followed. Strains are listed as carrying the *unc-119(ed3)* allele, but the genotype at this locus was not confirmed.

**Table S2.**
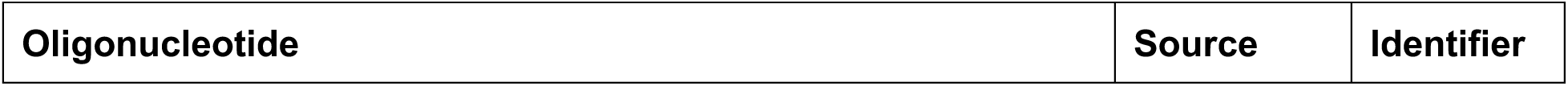
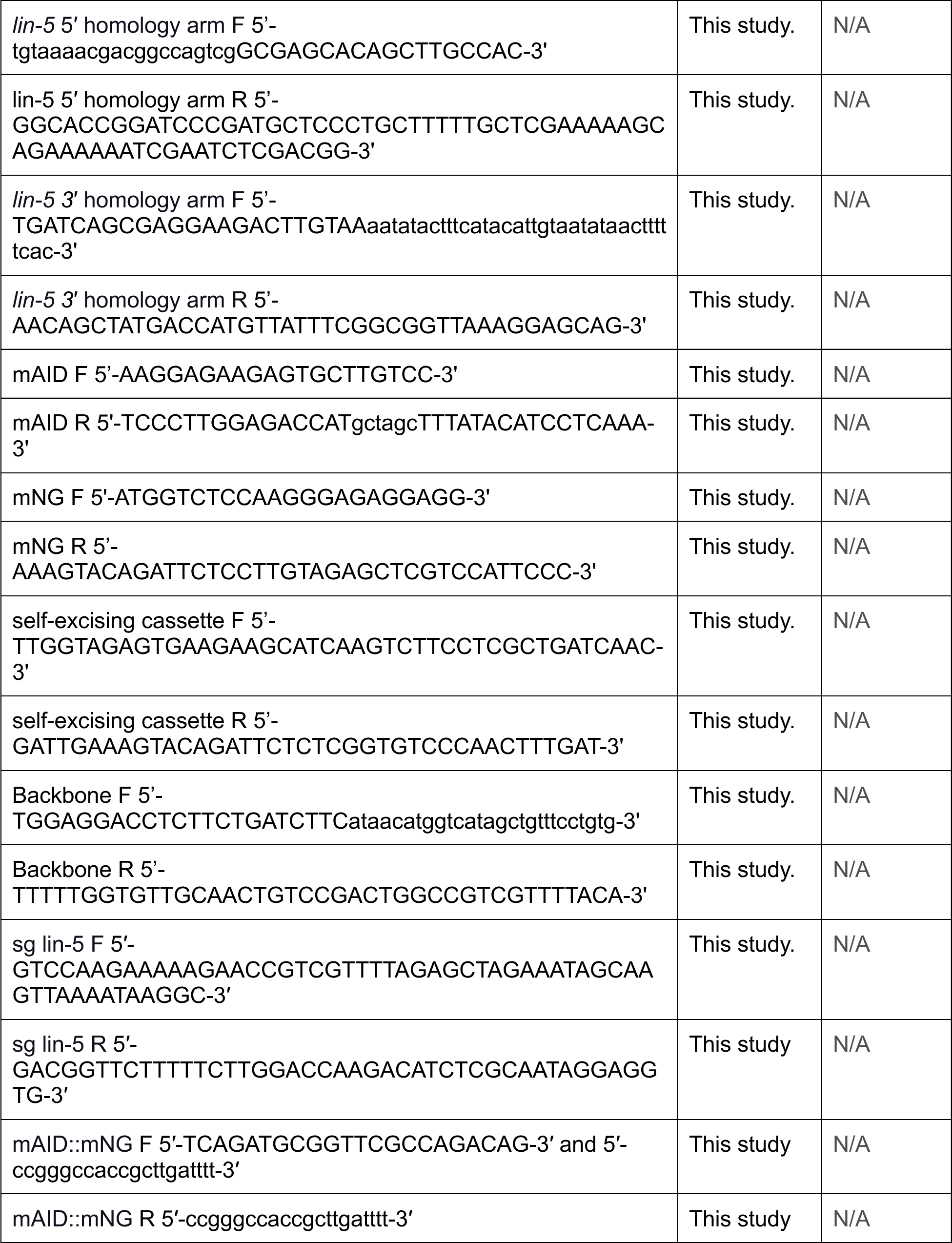
List of primers used in the generation of LIN-5::mAID::mNG.

**Table S3.** Imaging parameters. Imaging parameters are listed by experiment with relevant figures indicated. Z-step for all experiments is 0.5 µm. For all experiments done using the Zeiss Cell Observer SD a binning of 3x3 was used. For the experiments done using the Nikon CSU-X1 no binning was used. * = For all experiments done using the Zeiss Cell Observer SD a quad pass 466/523/600/677 filter was used. SV = Supplementary Video. ST = Single timepoint. SD = Spinning disk.

| Experiment | Microscope | Duration (min) | Z-slices | Frame rate (sec) | Laser (%) |  | Exposure (ms) |  | Filters | Objective | Figures |
| --- | --- | --- | --- | --- | --- | --- | --- | --- | --- | --- | --- |
|  |  |  |  |  | 488 | 561<br>568 | 488 | 561<br>568 |  |  |  |
| Dual channel imaging of mCH::TBB-2 and mNG::ANI-1 | Zeiss Cell observer SD | Up to 40 | 28 to 60 | 30 to 60 | 10<br>5 | 10 | 200 | 200 | * | 63x | All 1<br>All S1<br>S2C<br>SV1 |
| Dual channel imaging of mCH::TBB-2 and mNG::ANI-1 (+Brightfield (1<sup>st</sup> timepoint middle z-slice only)) | Nikon CSU-X1 | Up to 40 | 61 | 30 to 90 | 5 | 5 | 80 | 80 | 59004-FITC/<br>TRITC | 40x | 1G-I<br>S2D |
| Dual channel imaging of TagRFP::PH, mNG::ANI-1 and GFP::TBB-2 | Zeiss Cell observer SD | Up to 40 | 48 to 62 | 45 to 60 | 10 | 10 | 200 | 200 | * | 63x | 2B-H<br>5F-G |
| Dual channel imaging of TagRFP::PH, mNG::ANI-1 and GFP::TBB-2 | Nikon CSU-X1 | Up to 40 | 61 to 83 | 30 to 75 | 4 | 2 | 80 | 80 | 59004-FITC/<br>TRITC | 60x | 2BH<br>5F-G |
| Dual channel imaging of TagRFP::PH, UNC-59::GFP and GFP::TBB-2 | Nikon CSU-X1 | Up to 40 | 61 to 83 | 30 to 75 | 4 | 2 | 80 | 80 | 59004-FITC/<br>TRITC | 60x | All 2<br>5F-G |
| Single channel imaging of mCH::TBB-2 and TIR-1::mRuby | Nikon CSU-X1 | Up to 40 | 61 | 45 to 55 | - | 5 | - | 80 | 59004-FITC/<br>TRITC | 60x | 3C-F<br>S3A-C<br>5I-K<br>S5A |
| Dual channel imaging of mCH::TBB-2 | Nikon CSU-X1 | ST | 61 | - | 5 | 5 | 100 | 100 | 59004-FITC/TRITC | 60x | 3B |
| Dual channel imaging of DHC-1::AID::GFP, mCH::TBB-2 and TIR-1::mRuby | Nikon CSU-X1 | Up to 40 | 61 | 55 to 80 | 5 | 5 | 80 | 80 | 59004-FITC/TRITC | 60x | 3A-F<br>S3A-C<br>5I-K<br>S5A-B |
| Dual channel imaging of mCH::TBB-2 | Nikon CSU-X1 | ST | 61 | - | 15 | 5 | 200 | 200 | EM525/50<br>EM605/55 | 40x | 3B |
| Dual channel imaging of LIN-5::mAID::mNG, mCH::TBB-2 and TIR-1::mRuby | Nikon CSU-X1 | Up to 40 min | 61 | 80 to 145 | 15 | 5 | 200<br>1<sup>st</sup> time point only | 200 | EM525/50<br>EM605/55 | 40x | 3A-F<br>S3A-C<br>5I-K<br>S5A |
| Dual channel imaging of DHC-1::AID::GFP, TagRFP::PH, mCH::TBB-2 and TIR-1::mRuby | Nikon CSU-X1 | ST | 51 | - | - | 2 | - | 100 | 59004-FITC/TRITC | 40x | 3G |
| Dual channel imaging of DHC-1::AID::GFP, mKate2::HIM-4 mCH::TBB-2 and TIR-1::mRuby | Nikon CSU-X1 | ST | 61 | - | 5<br>10 | 10<br>15 | 100<br>200 | 100<br>200 | 59004-FITC/TRITC<br>EM700/75 | 40x | 3H |
| Dual channel imaging of LIN-5::mAID::mNG and mKate2::ANI-1 | Nikon CSU-X1 | ST | 61 | - | 30<br>20 | 20 | 100 | 100 | EM525/50<br>EM605/55 | 40x | S3D |
| Dual channel imaging of LIN-5::mNG and mCh::PH | Nikon CSU-X1 | Up to 40 min | 61 | 60 to 100 | 5 | 5 | 80<br>200 | 80<br>200 | 59004-FITC/TRITC | 60x | All 4<br>All S4<br>SV2-4 |
| Dual channel imaging of GFP::TBB-2 and mCh::HIS-11 | Zeiss Cell observer SD | Up to 40 min | 38 | 30 | 15 | 10 | 200 | 200 | * | 63x | 5C-E |
| Dual channel imaging of GFP::TBB-2 and mCh::HIS-11 | Nikon CSU-X1 | Up to 40 min | 27 | 30 | 3 | 10 | 200 | 200 | 59004-FITC/TRITC | 40x | 5E |
| Dual channel imaging of | Nikon CSU-X1 | ST | 71 | - | 5 | 30 | 300 | 300 | 59004-FITC/TRITC | 40x | 5A-B |
| mCH::HIS-58 and TBG-1::GFP |  |  |  |  |  |  |  |  |  |  |  |
| Single channel imaging of LAM-2::mKate2 + Brightfield | Nikon CSU-X1 | ST | 91 | - | - | 30 | - | 80 | 59004-FITC/TRITC | 40x | S2B |
| Dual channel imaging of Plag-2: mCherry LIM-7::GFP + Brightfield | Nikon CSU-X1 | ST | 113 | - | 5 | 5 | - | 80 | 59004-FITC/TRITC | 40x | S2A |

## Supplementary video legends

Video S1 was acquired on a Cell Observer SD spinning disc confocal (Zeiss; Yokogawa) using an AxioCam 506 Mono camera (Zeiss), with 3x3 binning, and a 63x/1.4 NA Plan Apochromat DIC oil immersion objective (Zeiss) in Zen software (Zeiss). mNG::ANI-1 is shown in cyan and mCH::β-tubulin is shown in red. Images were acquired every 37.57 seconds. Maximum intensity projections of 21 z-slices (0.5µm sectioning) centered on the mitotic spindle are shown. Video S2, S3 and S4 were acquired on a Nikon CSU-X1 spinning disk confocal microscope (Nikon TI2-E inverted microscope with a Yokogawa CSU-X1 confocal scanner, controlled by NIS-Elements software, using a Nikon Plan Apo Lambda 60x/1.4 NA Oil immersion objective. In Video S2 and S4, LIN-5::mNG is shown in inverted greyscale. In Video S3, LIN-5::mNG is shown in cyan and mCH::PH is shown in red. Images for S2, S3 and S4 were acquired every 67.37, 17.09 and 102.42 seconds, respectively. For Video S2, maximum intensity projections of 11 z-slices (0.5µm sectioning) centered on the cell’s basal surface are shown. For Video S3 and S4, maximum intensity projections of respectively 3 and 11 z-slices (0.5µm sectioning) centered on the mitotic spindle are shown. All videos were processed using Fiji. Briefly, images were bleach corrected using the histogram matching method and adjusted for brightness and contrast. Images were then converted to RGB and scaled 4 times to adjust for file compression during AVI conversion. TIFs were converted to AVI in ImageJ using JPEG compression and a 6 frames per second framerate. For all videos, scale bar = 5 μm.

**Video S1.** Centrosomes and rachis bridge dynamics during germ cell mitosis. Related to Figure 1.

**Video S2.** LIN-5::mNG at the germ cell basal cortex during prophase. Related to Figure 4. Cell in prophase is indicated by an arrow.

**Video S3.** LIN-5::mNG foci on the lateral cortex of a germ cell prior to anaphase. Related to Figure 4. The lateral cortex is indicated by an arrow.

**Video S4.** Centrosomal LIN-5::mNG is redistributed to the basolateral cortices at the end of mitosis. Related to Figure 4. Cell ending mitosis is indicated by an arrow.

